# Human Cognitive Ability and the P300 Event-Related Brain Potential: A Systematic Review and Meta-Analysis

**DOI:** 10.64898/2026.02.13.705728

**Authors:** Matthew J. Euler, Kirsten Hilger

**Affiliations:** Department of Psychology, University of Utah, 380 S 1530 E BEHS 502, Salt Lake City, Utah, USA; Institute of Psychology, Department of Psychology I, Würzburg University, Marcusstr. 9-11, Würzburg D-97070, Germany; Professorship for Differential Psychology, Personality Psychology and Psychological Diagnostic, Faculty of Human Sciences, Vinzenz Pallotti University, Pallottistraße 3, Vallendar D-56179, Germany

**Keywords:** Meta-analysis, intelligence, electroencephalography (EEG), event-related brain potential (ERP), P300

## Abstract

Human intelligence is essential to understand complex ideas, to engage in various forms of reasoning, to learn from experience, and to adapt to new situations by taking thought. The P300 event-related brain potential has been related to intelligence scores and thus represents as a promising biomarker of general cognitive ability. However, empirical results are characterized by enormous heterogeneity, and a quantitative assessment of this literature is lacking. This preregistered meta-analysis provides the first systematic overview of neuroscientific studies that have examined associations between general cognitive ability and the P300 in healthy adult participants. Out of 5641 articles screened, 49 studies with up to 381 effects were eligible for PRISMA-based meta-analytic comparison. Study quality was evaluated using a novel Study Design and Implementation Assessment Device, which we developed particularly for Individual Difference Research (DIAD-ID) and provide together with our analysis code as free online resources to support future meta-analyses. Confirming our hypotheses, a small but significant positive across-study association was observed for general cognitive ability and P300 amplitudes (*r* = .13; 95%-CI [.06, .19]), while a significant negative across-study association was revealed for P300 latencies (*r* = -.18; 95%-CI [-.24, -.13]). Study heterogeneity was substantial, and sub-analyses highlighted potential moderators such as type of the task during EEG recording. We discuss limitations, open questions, and provide concrete guidelines for future research on the neurobiological underpinnings of individual differences in cognitive ability.

## 1. Introduction

Human cognitive abilities are multifaceted and include the capacity to understand complex ideas, to engage in various forms of reasoning, to learn from experience, and to adapt effectively to the environment (Neisser et al., 1996). Established tests allow the calculation of so-called intelligence quotients that attempt to capture an individuals’ overall or general cognitive ability (GCA) within a single metric. Such scores have been shown to predict educational and occupational success (Schmidt & Hunter, 2004) as well as positive life outcomes including health and longevity (Deary et al., 2004). Understanding the neurobiological bases of individual differences in human cognitive abilities is therefore an important aim of ongoing research across multiple scientific disciplines.

Electroencephalography (EEG) is a non-invasive neuroscientific method to measure electric potentials on the scalp surface generated by the brain, or more specifically, by synchronized firing of cortical neurons (i.e., mostly parallelly arranged pyramidal neurons; Nidal & Malik, 2014). In contrast to other methods like fMRI, EEG has high temporal resolution and allows to assess brain activity changes in the range of milliseconds. EEG can be used to capture spontaneous neural activity as well as event-related brain potentials (ERPs), i.e., the average electrophysiological response to a frequently presented stimulus or behavior. As such, it provides an excellent means for relating individual differences in cognitive abilities to momentary aspects of neurophysiological activity.

The P300 ERP or P3 is defined as the third positive deflection to a specific consciously detected stimulus and typically peaks around 300 milliseconds after stimulus presentation (Luck, 2014). A distinctive property of the P3 is that it is much larger for infrequently occurring stimuli than for frequently occurring stimuli. This can be observed in so-called oddball paradigms, in which the rare oddball stimuli elicit a larger P3 than the frequent standard stimuli. The P3 is suggested to reflect some type of higher-level cognitive processes (Luck, 2012) and many studies have been conducted to test whether individual differences in cognitive abilities relate to P3 amplitudes, reflecting the strength of neural activation, or to P3 latencies, informing the speed of neural information processing (Hilger et al., 2022). Although medium-to-large negative associations between cognitive abilities and P3 latency have been reported in studies using oddball paradigms (e.g., Bazana & Stelmack, 2002; De Pascalis et al., 2008; Saville et al., 2016; Stelmack & Houlihan, 1995; Troche et al., 2009; Walhovd et al., 2005), potentially supporting the mental speed hypothesis of intelligence (Der & Deary, 2017), effect sizes differ enormously and contradictory findings have also been observed (Nieman et al., 2002; Polich et al., 1992; Wronka et al., 2013). Moreover, the relation between cognitive abilities and the P3 latency in other cognitive tasks like the Sternberg Short-Term Memory Scanning Task, the N-back task, or the Hick task presents an open question, as negative (e.g., Jungeblut et al, 2021; Saville et al., 2016; Schubert et al., 2017; Schubert et al., 2021), positive, or no relations have been found (Euler et al., 2017; Houlihan et al., 1998; Schubert et al., 2018; Troche et al., 2017). Finally, this heterogeneity also concerns investigations focused on P3 amplitudes. Again, positive (Russo et al., 2008; Varriale et al., 2018), negative (Wongupparaj et al., 2018; Ramchurn et al., 2014) as well as no associations (Euler et al., 2017) with cognitive differences have been observed, thereby preventing firm conclusions.

The above outlined heterogeneity not only pertains to the size and direction of reported effects, but also to various characteristics of study design. Beyond different tasks employed during EEG assessment, studies differ also in their measurement of individual differences in cognitive abilities, the definition of the P3 (e.g., in which time window), and characteristics study samples (e.g., only students vs. representative community samples). Unfortunately, although the P3 presents a promising candidate marker of human cognitive abilities, a systematic evaluation of this literature has not yet been conducted.

This is even more problematic, as studies in behavioral and psychological sciences are in a “replication crisis” (Lakens, 2015). Many of them suffer from a lack of transparency and the use of questionable research methods, preventing replication (Open Science Collaboration, 2015). Open, transparent, and reproducible research practices have been proposed as a solution to this problem (e.g., preregistration) together with systematic reviews and meta-analyses (e.g., Arslan, 2019; Polanin et al., 2020). Systematic reviews and meta-analyses provide more valid evidence than individual studies alone (Müller et al., 2018) and, most critically, allow identification of potential publication bias, to estimate the risk of bias in the study design, and ultimately to derive best research practices (Polanin et a., 2021).

Given that the P3 is among the most-studied ERPs, a meta-analytic assessment of its relation to human cognitive abilities is arguably past due. Further clarifying the presence, size, and direction of its relation to individual cognitive abilities is important for guiding future electrophysiological research on the nature and correlates of human intelligence. An improved understanding of the factors that moderate those relationships should not only enhance theory-development on the neural basis of cognitive differences, but will also inform clinical and other translational applications of the P3 where individual differences in cognitive abilities are relevant (e.g., pre-clinical detection of dementia risk; Paitel et al., 2021; physiological monitoring of cognitive load; Ghani et al., 2020; Lean, & Shan, 2012).

Here, we provide a systematic evaluation and meta-analytical comparison of all studies published before 2024, investigating associations between individual differences in cognitive abilities and different features of the P3 in healthy adult participants. Given the hierarchical structure of cognition—the fact that different mental abilities are highly correlated, and individual differences in mental abilities are well-captured by measures of general cognitive ability or intelligence (Carroll, 1993; Haier, 2023)—our meta-analysis focuses on GCA as the cognitive construct of interest. However, we extracted information about constituent abilities where this information was available and systematically evaluated potential moderators as well as study design characteristics to derive insights into how these may affect study results. Systematic variation among studies, important shortcomings, potential publication bias, and pressing challenges in this field of research were also considered with the ultimate goal of providing useful recommendations, guidelines and resources for future research.

## 2. Hypotheses

In addition to addressing the core claims that P3 amplitude and latency are associated with human cognitive abilities, we also propose hypotheses regarding potential moderators. Given their established relevance in the neuroscience of cognitive ability (Euler & Schubert, 2021; Larson et al., 1995; Neubauer & Fink, 2009), we focused on task type and task difficulty. Therefore, the following hypotheses were preregistered on the Open Science Framework: https://osf.io/2xst4.

**H1.** P3 amplitudes are significantly positively associated with general cognitive ability. **H1a.** The sign of the association between P3 amplitude and GCA is moderated by task type. Specifically, we hypothesize that the relation is positive when the P3 is elicited by Oddball paradigms, and negative when the P3 is elicited by paradigms that require levels of cognitive control exceeding that of stimulus recognition (e.g., memory encoding and retrieval; see Wongupparaj et al., 2018; Wronka et al., 2013).

**H1b.** The magnitude of the association between P3 amplitude and GCA is moderated by task type. Specifically, the size of the association increases with greater task difficulty (Bazana & Stelmack, 2002; Sculthorpe et al., 2009).

**H2.** P3 latencies are significantly negatively associated with general cognitive ability. H2a. The magnitude of the association between P3 latency and GCA increases with greater task difficulty (e.g., Kapanci et al., 2019).

## 3. Methods

### 3.1 Preregistration

The protocol for this study was formally preregistered on the Open Science Framework: https://osf.io/2xst4. Please note that during literature search, it was necessary to deviate from our pre-registered search terms to accommodate the requirements of selected databases. Additionally, it was necessary to refine our exclusion criteria during the title/abstract screening and full text review stages to limit the number of irrelevant studies. Both deviations are described in more detail in the relevant sections below (3.2, 3.2.1, 3.2.2).

### 3.2 Literature Search and Study Selection

Studies published until June 2023 were retrieved and selected using the Preferred Reporting Items for Systematic Reviews and Meta-Analyses (PRISMA; Page et al., 2021). Test searches were conducted to refine keywords. We consulted the electronic databases APA PsychInfo, Google Scholar, ProQuest Theses and Dissertations Global, Pubmed, Scopus, Web of Science, bioRxiv, PsyArXiv, and ResearchGate with the search terms: (P3 OR P300 OR P3a OR P3b) AND (“cognitive ability” OR intel* OR “IQ” OR “general intelligence” or “fluid intelligence”) AND (ERP OR “event-related potential” OR EEG).

The following minor modifications had to be made to accommodate requirements of selected databases: First, Google Scholar does not allow batch export of search results, which necessitated the use of the Publish or Perish software (Harzing, 2007). We limited the number of results to 1000, which provided 862 records. Second, searching in ProQuest and Theses Global yielded several thousand studies (∼5500), many of which were irrelevant (concerning clinical conditions or child samples). Thus, that we further limited that search by the addition of: NOT (schiz* OR disorder OR disease OR depression OR bipolar OR mania OR children OR adolescents). We also limited the search in ProQuest to Doctoral Dissertations Only (January 1, 1907-May 1, 2023, English or German language only). Third, for bioRxiv, the original search terms had to be shortened due to exceeding the character limit of the search function. The following modification was used instead, returning 264 results: (P3 OR P300) AND (“cognitive ability” OR intel* OR “IQ” OR “general intelligence”) AND (ERP OR “event-related potential”). We uploaded the first 200 returned results ordered by Best Match. Fourth, since ResearchGate hosts many types of materials, we excluded books, data, presentations, and posters; and used the original terms to search for articles, literature reviews, theses, conference papers, and preprints. Fifth, as PsyArxiv and ResearchGate do not allow batch output, individual records from these databases were manually exported and uploaded to the review management software (see below) together with all other records for further processing. Finally, we identified additional 29 articles of possible relevance during title/abstract screening and full text review, which we also included.

#### 3.2.1 Title and Abstract Screening

Our search resulted in a total of 5641 articles, which were uploaded to the Covidence systematic review management software for screening https://www.covidence.org/. The software performs an automated de-duplication, which removed 1065 articles, leaving a total of 4523 for title and abstract screening (Figure 1). This was performed by two reviewers independently. Any studies fulfilling the following criteria were excluded: use of non-human subjects, focus on children or adolescents (< 18 years); primary focus on neurological, psychiatric, or medical conditions (e.g., cardiopulmonary disorders, endocrine/metabolic conditions, autoimmune disorders, etc.); task not defined well-enough to allow clear categorization (e.g., Oddball, memory encoding, memory retrieval, cognitive control, etc.); P300 not measured from scalp electrodes or measurement approaches being too idiosyncratic to allow coding of electrodes, measurement windows, etc.; article not available in English or German.

**Figure 1.**
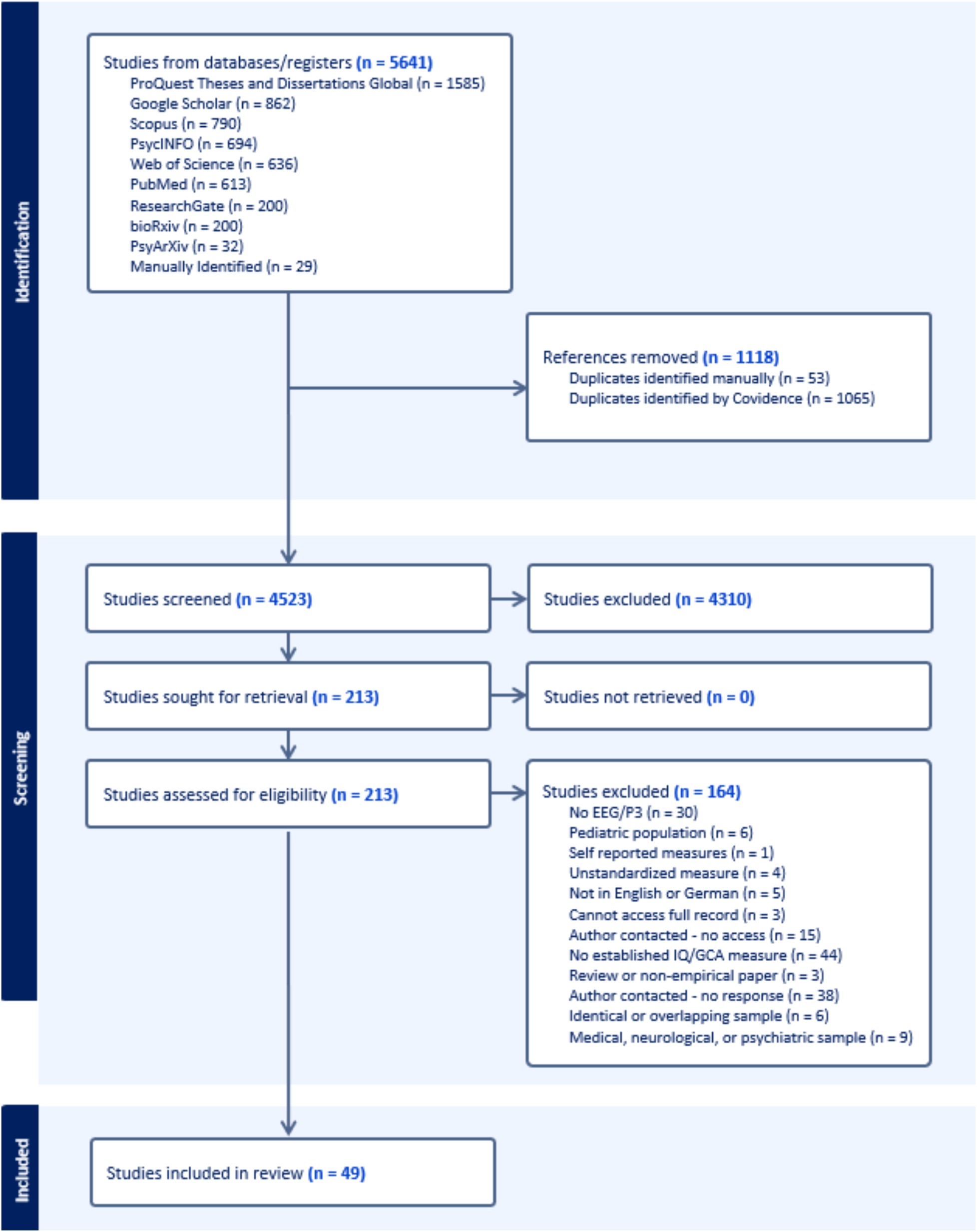
PRISMA flowchart of the study selection process. Of 4523 screened articles, 49 were included in the systematic review and quantitative meta-analysis.

After a brief period of initial screening, we decided to slightly deviate from these original pre-registered criteria to reduce the number of irrelevant articles going forward to full text review. Specifically, we added two criteria to also exclude studies where the “abstract makes no reference to individual differences in cognitive processes, tasks, or assessments” or studies that constitute “a review, meta-analysis, commentary, opinion, or other article type than an empirical study,” and began screening again. If the above criteria could not be ascertained from the abstract, the record was retained for further examination at the full text review stage. Intervention studies (e.g., cognitive training programs designed to increase cognitive abilities), were included if they collected both P3 and cognitive ability measures. Conflicts between both raters were discussed and solved. After screening, 213 studies met criteria for full-text review.

#### 3.2.2 Full-Text Review

At the full-text review stage, we included any study that passed our previous criteria and that reported a correlation between the P3 amplitude or latency and one or more measures of general cognitive ability (GCA). This step was performed by two raters independently. Due to the need to define GCA more precisely, we added to the preregistered exclusion criteria a criterion to exclude studies that did not use standardized, published, and established tests of GCA (e.g., the Wechsler Adult Intelligence Scales and subtests therein, Wechsler, 1955; the Ravens progressive matrices, Raven, 2003; Cattell Culture Fair test, etc., Cattell, 1973), or studies that focused on measures primarily used for neuropsychological assessment and diagnosis (word-list learning tasks, the Wisconsin Card Sorting Task; Sherman et al., 2023). Given their high correlation with and established use as proxy measures of GCA, studies using validated tests of irregular word-reading, such as the National Adult Reading Test (Bright et al., 2018; Nelson & Willison, 1991) were included.

For studies that did not report an association between the P3 amplitude or latency and a GCA measure but appeared to have collected relevant data, we contacted the authors in a standardized procedure (see preregistration). Note that we refrained from conducting any raw data processing ourselves (e.g., calculating P3 variables from EEG raw data), but included studies where the authors provided GCA and P3 amplitudes/latencies and we only had to calculate the correlations. Again, conflicts between both raters were discussed and solved. Finally, a total of 49 studies remained for inclusion in the systematic review and meta-analyses.

### 3.3 Data Extraction and Variables of Interest

The following information was extracted from each study:

- <u>P3-GCA Association</u>: Any correlation between a P3 amplitude/latency measure and a GCA assessment reported in the text of the original studies, provided by the original study authors or calculated by the authors of the current meta-analysis, representing the main variable of interest.
- <u>Basic Study Information</u>: Author names, publication year, DOI, and whether the study was peer-reviewed or gray literature.
- <u>P3: Amplitude/latency measure (peak, mean, etc.)</u>: amplitude/latency time window (start and end time); and descriptive statistics (mean, standard deviation, minimum, maximum) for P3 amplitudes/latencies at electrodes Fz, Cz, Pz, and “Other”.
- <u>GCA</u>: Name of the measure, number of (sub)tests administered, and total testing time. We also extracted the specific domain intended to be assessed by a given measure (e.g., Ravens matrices: fluid intelligence, vocabulary tasks: crystallized intelligence, etc.). Descriptive statistics for Wechsler IQ scores (mean, standard deviation, minimum, maximum), were also extracted if reported.
- <u>EEG Recording</u>: Sampling rate; online and offline reference; online and offline high-pass, low-pass, and notch filters.
- <u>Task Characteristics:</u> The name of the task given by the original study authors and task type as categorized by the authors of this meta-analysis (Oddball, Attention or Cognitive Control, Working Memory, and Reaction Time/Chronometric). Any tasks that could not be readily categorized were grouped into a single category “Other.”
- <u>Task Difficulty</u>: For studies that reported multiple P3-GCA associations based on different tasks (McGarry-Roberts et al., 1992) and/or conditions (e.g., Frischkorn, 2019; Houlihan, 1994), we inferred their related difficulty level by examining accuracy and/or reaction time differences, and by considering statements in the text denoting aspects like e.g., “complexity,” “task demands,” “switching costs,” etc. For studies reporting associations for active vs. passive or rare vs. standard oddball conditions, we also coded these as high and low difficulty, respectively (e.g., Russo et al., 2008). Cases where this information was ambiguous or unavailable were not coded.
- <u>Sample Information</u>: Description of the study sample given by the original study authors (e.g., “college students,” “healthy adults,” etc.); presence of subjects below 18 years old; descriptive statistics for participants age (minimum, maximum, mean, median, standard deviation); percent female, male, and non-binary participants; percent right-handed; study location; educational and occupational demographics; and the final sample size. Whereas most demographic information describes the whole sample, final sample sizes (after any exclusions took place) were recorded for each P3-GCA association separately. Finally, since some studies contain multiple sub-samples (due to sub-studies or sub-groups within a single study) P3-GCA associations were coded on a *per sample* basis wherever applicable and sample rather than study was used as the primary variable for modelling between-study heterogeneity.

#### Exceptional cases and caveats

Note that for studies that reported P3 amplitude, latency, or GCA values by low and high-ability groups separately, P3-GCA associations were pooled manually for further analyses.

- First, a single study examined the effect of signal processing choices on P3-GCA associations (Schubert et al., 2023), increasing the potential number of effects for analysis 12-fold. All variants were retained in the systematic review and descriptive analyses. However, to avoid over-weighting this study in the quantitative meta-analysis, only the effects resulting from the most commonly-accepted variants were included in our statistical model (average reference, 16 Hz low-pass filter; Luck, 2014).
- Second, although fractional area latency has been recommended as being the more reliable measure for latencies compared to peak latency (Luck, 2014), given the much more frequent use of peak latency in the available empirical evidence, we chose peak latency to calculate the primary meta-analytical latency effects.
- Third, for demographic variables, many studies only report information for the initially enrolled study sample rather than for sub-samples and typically not for the final sample that was subject to correlational analyses (after any exclusion took place). Therefore, we considered the demographic information that was available (typically of the enrolled sample) in the Results section of our systematic Review.

### 3.4 Study Design Quality Assessment

To systematically and objectively assess study quality and potential biases in study design, all studies were rated with our newly developed Study Design and Implementation Assessment Device for Individual Difference research (DIAD-ID), which is based on the Study Design and Implementation Assessment Device (DIAD) by Valentine and Cooper (2008). The DIAD allows a tailored evaluation of potential bias in research studies and can be applied across various fields of research, inclusively non-clinical studies (Bernard et al., 2014; Linhardt et al., 2022; Pfeiffer et al., 2024). Our DIAD-ID presents a valuable modification of this system and can easily be adapted to any neuroscientific research on individual differences. First, contextual questions must be specified to adapt the rating system to a specific field of research (context). We defined 12 questions, whose final answers constitute the main goal of the algorithm, and that can be grouped into (A) Construct validity: Fit between concepts and operations, (B) Construct validity: Fit between Concepts and outcomes, (C) Construct validity: Causal Inference, (D) External validity: General, (E) External validity: Sampling, (F), External validity: Representativeness of effect testing, (G) Statistical validity: Effect size estimation, (H) Statistical validity: Effect calculation, and (I) Statistical validity: Reporting. These 12 questions were rated through 36 more specific design and implementation questions, which were newly developed and present the input for the algorithm. The DIAD-ID, inclusively all questions and example ratings, is ready to be used in future reviews and can be accessed online: https://osf.io/j9g7q/.

### 3.5 Examination of Outliers

Potential outliers were identified at the individual study level, based on the joint criterion of studentized residuals ± 1.96 (Viechtbauer & Cheung, 2010), and a Cook’s distance value larger than the median value plus six times the interquartile range, as calculated separately for the associations between GCA and P300 amplitude as well as for the association between GCA and P300 latency (Pfeiffer et al., 2024). For any study identified by these criteria, we conducted sensitivity analyses to understand their potential influence on the meta-analytical across-study association, its significance, and on the heterogeneity metrics.

### 3.6 Meta-Analytic Synthesis of Results

All analyses were conducted in R (version 4.4.2; R Core Team, 2024) using the following packages: tidyverse (Wickham et al., 2019), RColorBrewer (Neuwirth, 2022), moments (Komsta & Novomestky, 2022), pander (Daróczi & Tsegelskyi, 2025), psych (Revelle, 2025), stringr (Wickham, 2025), forestplot (Gordon & Lumley, 2025), and metafor (Viechtbauer, 2010). The effect sizes for P3-GCA associations (correlations) from the original studies were converted via the Fisher *z*-transformation and weighted by sample size. The main hypotheses concerning associations between GCA and P3 amplitudes and latencies (H1 and H2) were tested with intercept-only meta-analytical models with sample as a random effect. This allowed for estimating the average across-study effect size for amplitude and latency associations while accounting for between-study heterogeneity. To test the sub-hypothesis that Task Type moderates the association between GCA and P3 amplitude (H1a), Task Type was entered as a categorical fixed effect with five levels (“Oddball”, “Attention/Cognitive Control”, “Working Memory”, “Chronometric”, and “Other”), with “Oddball” as the reference category, while sample was again included as random effect. We fit analogous models to test the sub-hypotheses that Task Difficulty moderates the association between GCA and P3 amplitude (H1b) and P3 latencies (H2). Difficulty was entered as a categorical fixed effect with three levels (“Low”, “Medium”, “High”), with Low as the reference category, and sample was again modelled as random effect. All models were fit using the Restricted Maximum Likelihood estimator (default in metafor) after ensuring that statistical assumptions were met.

### 3.7 Publication Bias

Publication bias was evaluated at a sample-level, while multiple effects within a sample were aggregated after Fisher’s *z* transformation and fixed-effect inverse-variance weighting. Funnel plots were inspected graphically, and statistical asymmetry was tested with Egger’s regression and the rank-correlation test (Begg and Mazumdar, 1994) applied to a univariate random-effects model. Where asymmetry was suggested, trim-and-fill imputation was used to estimate an adjusted effect size. As file-drawer analyses, we computed the generalized fail-safe *N* (implemented in *metafor* for random-effects models), which estimates the number of null studies required to render the overall effect non-significant at alpha = .05.

### 3.8 Data and Code Availability Statement

All software and packages used in the current study are open source. All analysis code used to conduct the meta-analyses and to create figures included in this paper is available on OSF: https://osf.io/j9g7q/. Note that we also share our extraction database and coding schemata for full transparency as well as the further developed DIAD (the DIAD-ID) that now can be easily transferred to any meta-analyses on brain-behavior relationships: https://osf.io/j9g7q/.

## 4. Results

### 4.1. Systematic Review

#### 4.1.1. Study Sample Characteristics

Forty-nine studies met criteria for inclusion in the quantitative meta-analyses, comprising 55 unique samples, *N* = 3107 participants, and 1612 total P3-GCA associations (across samples, experimental conditions, and electrodes, and including both amplitudes and latencies). Studies varied in the number of reported associations and sample sizes ranged widely (*M* = 56.22 participants, *SD* = 37.18, *Range*: 8-159; Table 2), with most falling far below recent recommendations for achieving adequate power in individual differences research (DeYoung et al., 2022, 2025; Schönbrodt & Perugini, 2013). Study samples typically consisted of young, female, right-handed participants, though 13 studies examined adults spanning a wider range of age (e.g., Walhovd & Fjell, 2003). Two studies focused exclusively on older adults (> 60; Table 2; e.g., Spironelli et al., 2020). Although studies with a primary focus on pediatric samples were excluded, five studies included a small number of older adolescents within their primary adult sample (e.g., Varriale et al., 2018). Educational levels and other sample demographic characteristics were reported infrequently (Table 2).

**Table 1.**
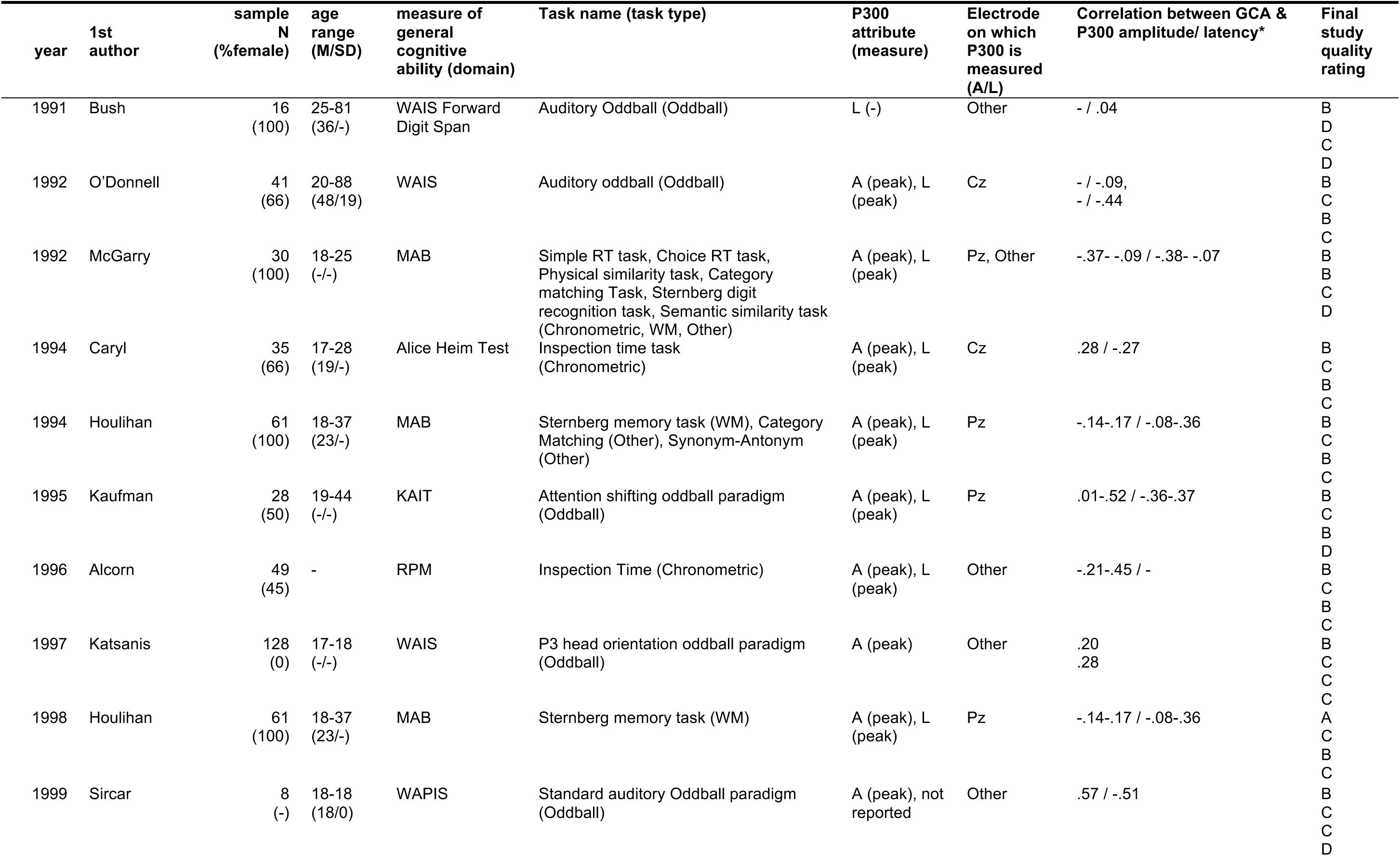
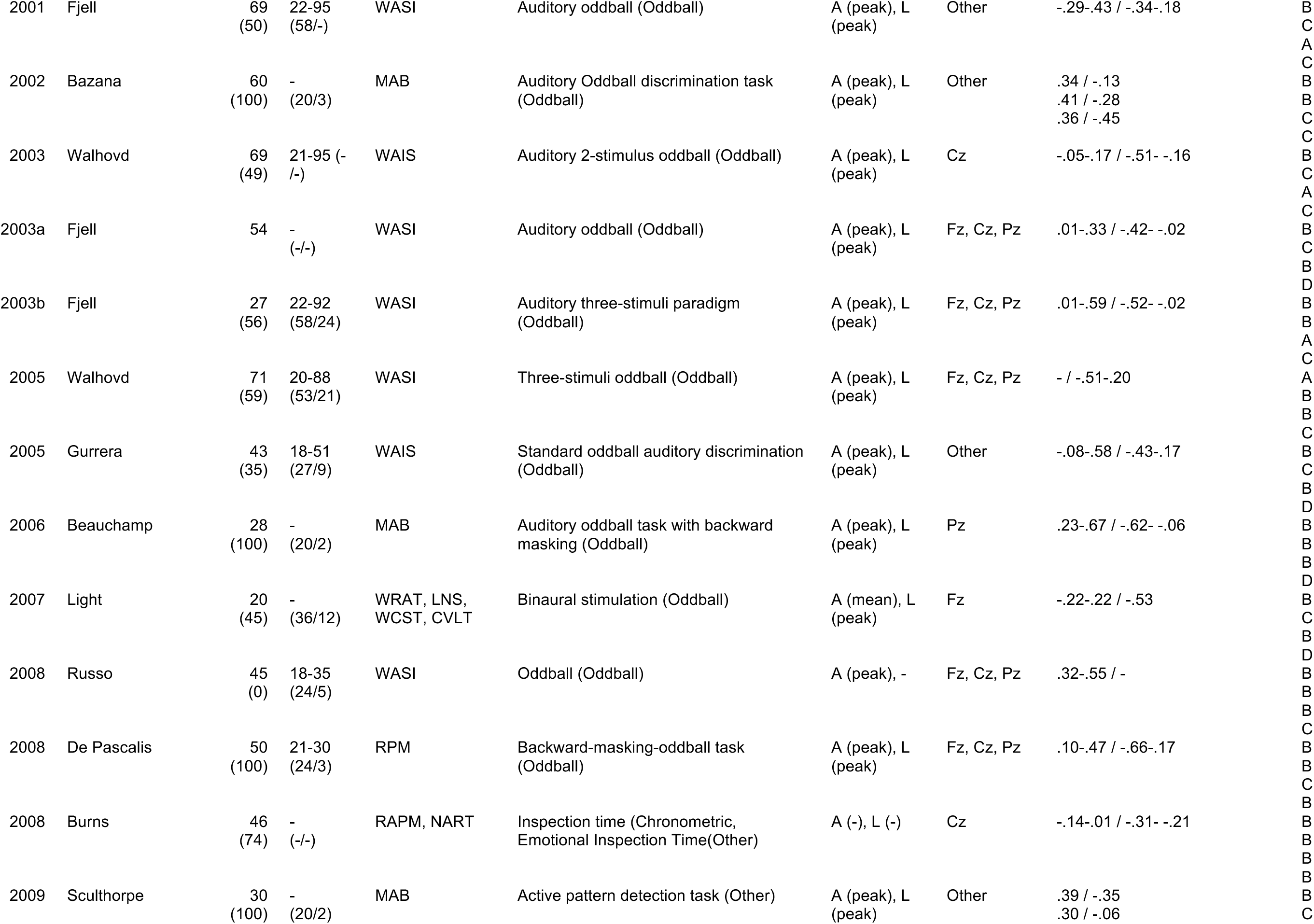
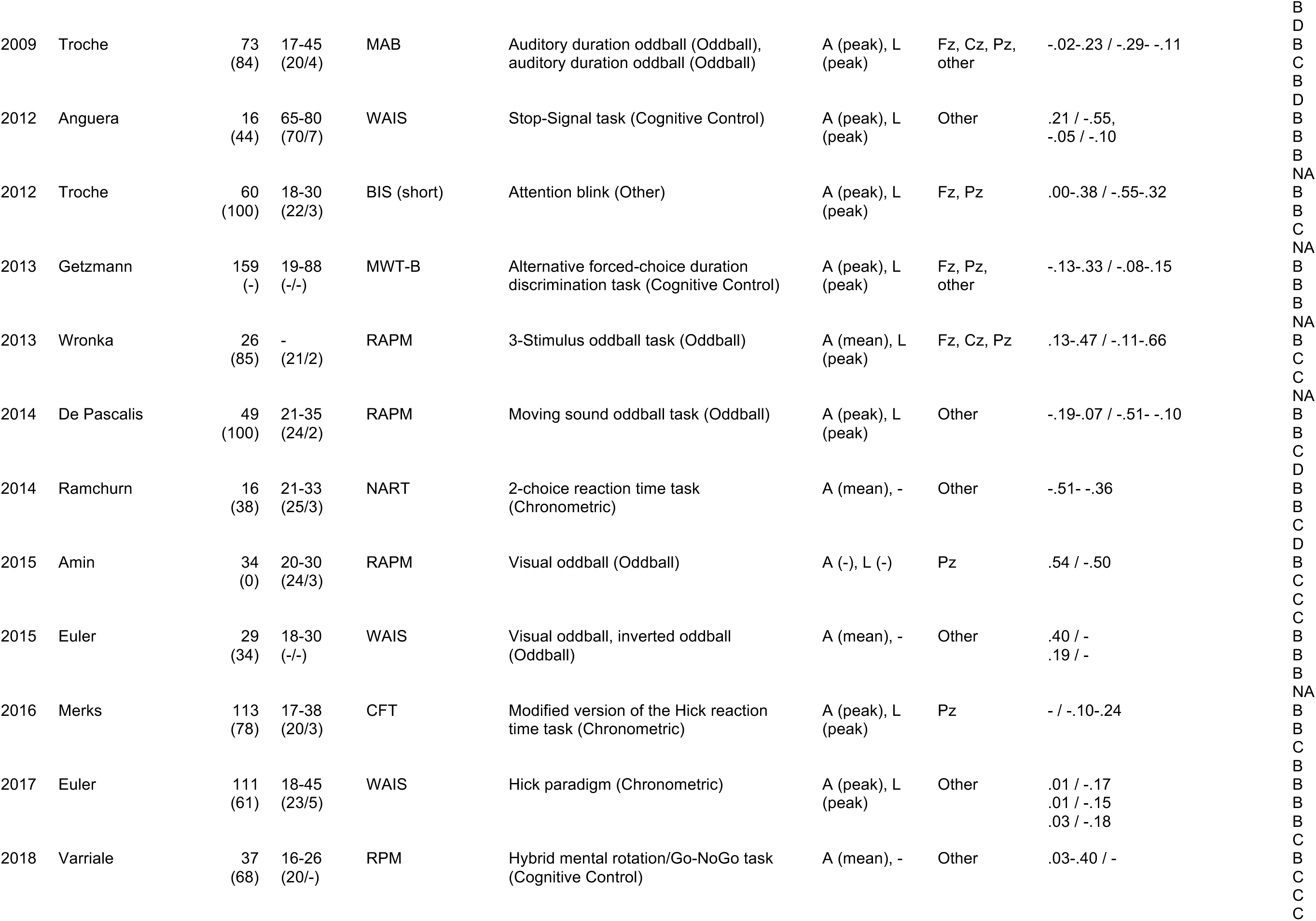
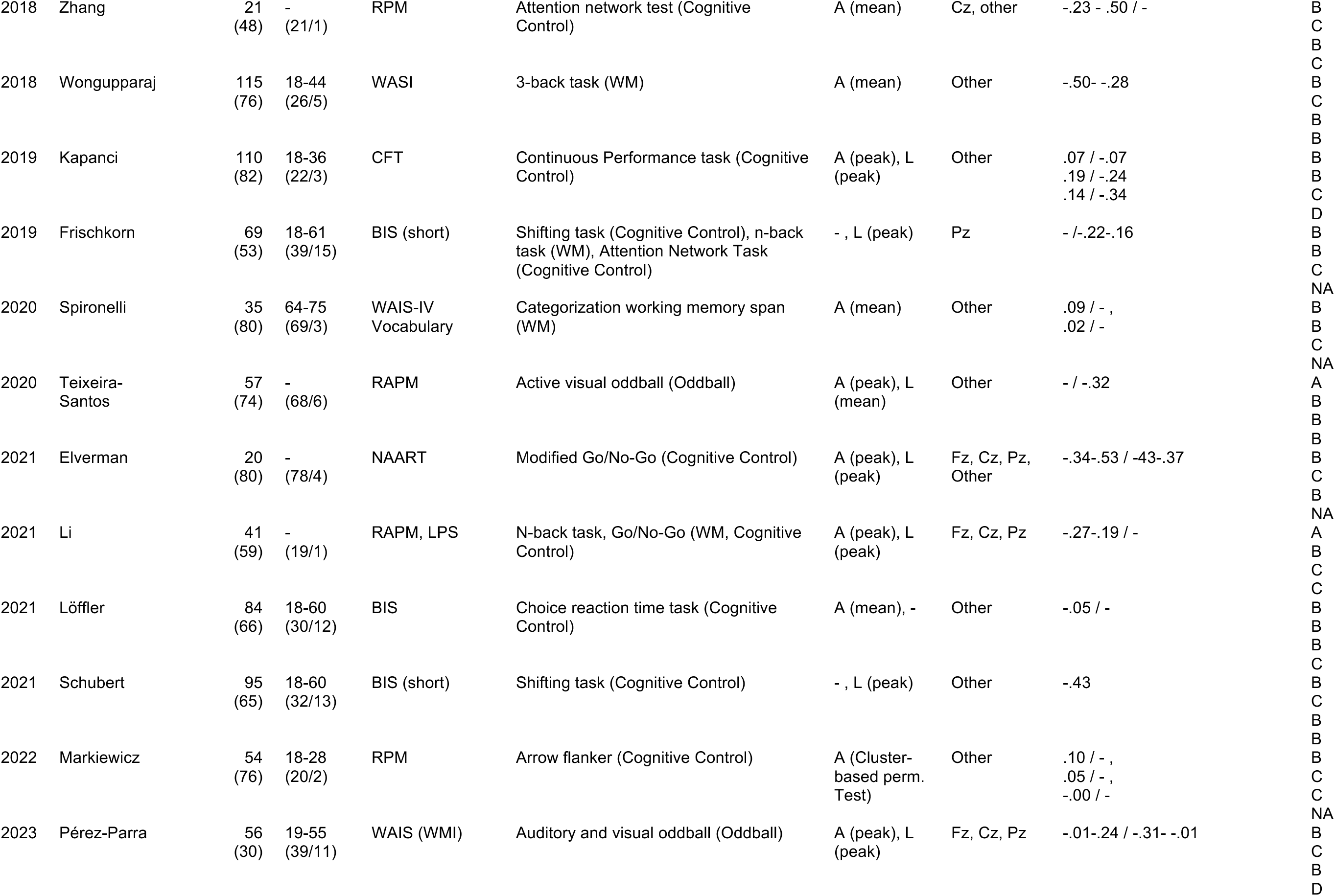
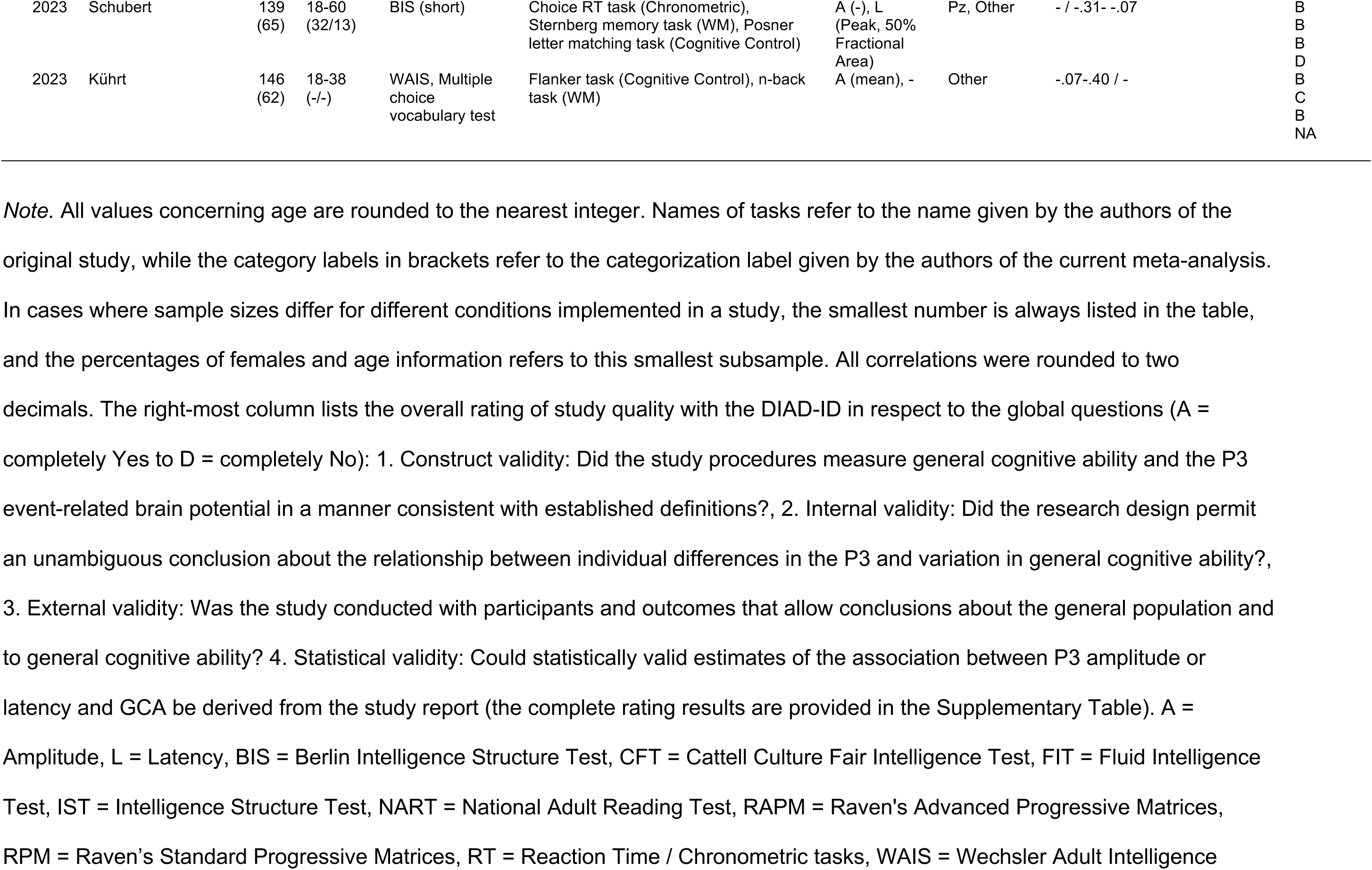
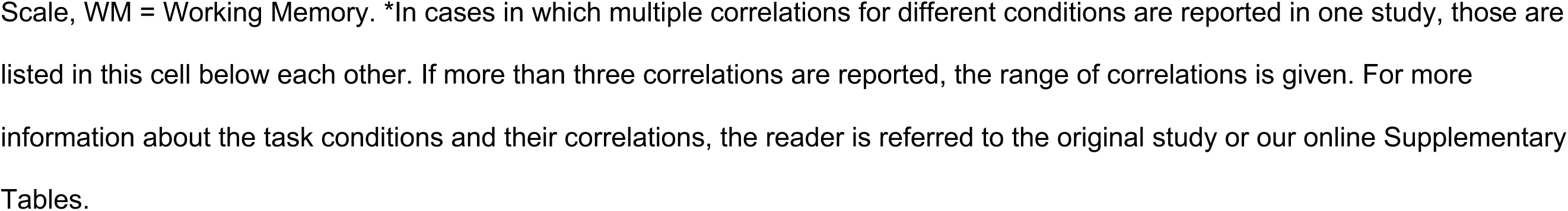
Studies Included in the Meta-Analyses in Chronological and Alphabetical Order.

**Table 2.** Demographic characteristics of the meta-analytical sample and reporting frequencies.

| Variable | Mean | Median | SD | Min | Max | Skew | Reporting Frequencies |  |
| --- | --- | --- | --- | --- | --- | --- | --- | --- |
|  |  |  |  |  |  |  | Studies | Samples |
| Sample Size | 56.22 | 46 | 37.18 | 8 | 159 | 1.10 | 49 | 55 |
| Mean Age | 32.36 | 23.7 | 17.79 | 18.4 | 79.67 | 1.48 | 39 | 45 |
| SD Age | 6.04 | 3.19 | 5.93 | 0.26 | 24 | 1.49 | 33 | 39 |
| Age Youngest | 21.23 | 18 | 10.48 | 16 | 65 | 3.83 | 37 | 38 |
| Age Oldest | 50.72 | 44 | 23.40 | 18 | 95 | 0.69 | 36 | 37 |
| % Female | 66.30 | 66 | 26.62 | 0 | 100 | -0.61 | 46 | 52 |
| % Right-Handed | 97.94 | 100 | 3.77 | 88 | 100 | -1.58 | 22 | 25 |
| Mean Edu. | 14.39 | 14.38 | 1.21 | 11.06 | 16 | -1.44 | 9 | 13 |
| SD Edu. | 2.03 | 2.2 | 0.81 | 0.59 | 3.1 | -0.52 | 9 | 13 |
| Min Edu. | 10.5 | 10.5 | 0.71 | 10 | 11 | 0 | 2 | 2 |
| Max Edu | 18 | 18 | 0 | 18 | 18 | — | 2 | 2 |
| % College Students | 92.21 | 100 | 22.82 | 7 | 100 | -3.11 | 16 | 19 |
| Sample IQ Mean | 110.35 | 111.28 | 6.70 | 94.58 | 121.6 | -0.75 | 21 | 24 |
| Sample IQ SD | 11.30 | 10.89 | 4.33 | 3.59 | 23.3 | 0.95 | 19 | 22 |
| Sample Minimum IQ | 83.33 | 85.5 | 8.97 | 61 | 96 | -1.21 | 11 | 12 |
| Sample Maximum IQ | 137.71 | 137 | 4.54 | 130 | 147 | 0.67 | 11 | 12 |

*Note.* Values are rounded to the nearest decimal. Studies refers to the number of studies reporting the variable of interest. Samples refers to the number of samples for which that variable is reported. Educational statistics are reported in years. % = percentage. IQ = intelligence quotient.

#### 4.1.2 Cognitive Ability Measurements

Associations with P3 amplitudes/latencies were examined approximately equally often for fluid intelligence (30.6%) and GCA/general intelligence (30.6%), followed by crystallized intelligence (18.06%), working memory (11.1%), visuospatial skills (5.6%), and processing speed (4.2%; percentages refer to GCA domains across samples, Figure 2A). On average, each sample was administered 4.46 tests to assess cognitive ability, with a median of 1 (*SD* = 4.78, *Range* = 1 - 15). These tests covered roughly two abilities (e.g., fluid and crystallized intelligence; *Mean* = 1.89, *Median* = 1, *SD* = 1.76, *Range* = 1-7). Twenty-one studies (43.6% of samples) reported mean IQ values in the standardized Wechsler metric. Averaging across these, the mean IQ of participants in our meta-analysis was 110.3 (*SD* = 6.7, *Range* = 94.6-121.6, *Skew* = -0.75) with an average *SD* of 11.30 (*Range of SDs*= 3.59 - 23.3). Study samples varied in their IQ ranges with an average lowest IQ value of 83.33 (*Range of Minimum Values* = 61 - 96), and an average highest IQ value of 137.70 (*Range of Maximum Values* = 130 - 147). Therefore, the study samples included in our meta-analysis were characterized by slightly above-average IQs, less variability around the mean, and a skew toward higher-ability individuals compared to the population (see Table 2).

**Figure 2.**
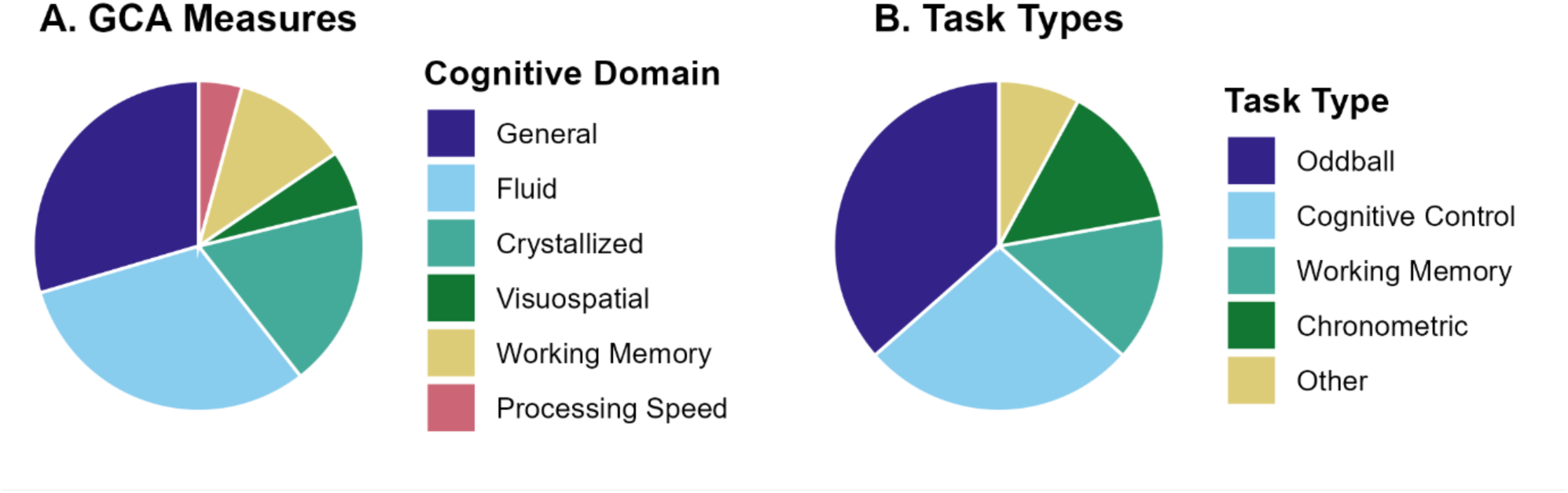
Most studies examined fluid intelligence and elicited the P3 using oddball tasks. (A) Proportions of general cognitive ability domains examined in meta-analytically included samples. (B) Proportions of experimental task types administered during EEG recordings in the meta-analytically included samples.

#### 4.1.3. Study Task Characteristics

P3-GCA associations were reported most frequently for oddball tasks (37.5%; e.g., Walhovd & Fjell, 2003), followed by cognitive control tasks (26.6%; e.g., Flanker, Go/No-Go; Markiewicz et al, 2021; Elverman et al., 2021), working memory (14.1%; e.g., N-back, Sternberg memory scanning; Li et al. 2021; Schubert et al., 2023) and chronometric tasks (14.1%; e.g., inspection time, Alcorn & Morris, 1996; Hick paradigm, Merks, 2016) with 7.8% classified as “other” (percentages refer to task types across samples; Figure 2B). These percentages differed slightly when calculated across all available associations (*m* = 1612), with oddball-based measures representing 41.7% of associations, followed by cognitive control (20.3%), working memory (16.4%), chronometric (15.6%) and “other” (6.0%).

#### 4.1.4 EEG Methodology and Signal Processing

With respect to recording parameters and preprocessing steps, 42 of the 49 included studies reported data on digital sampling rates (85.7%), with a modal value of 500 Hertz (Hz; *Range*: 100 – 1024). Thirty (61.2%) and 35 (71.4%) studies reported information on online high- or low-pass filters, respectively, while 15 (30.6%) and 28 studies (57.1%) reported information on offline application of high- or low-pass filters. The modal value for both online and offline high-pass filters was 0.1 Hz (*Range:* 0.01 – 5 Hz, online; *Range*: 0.01- 3 Hz, offline), while the modal value was 100 Hz for online low-pass filters (*Range*: 10 – 200 Hz), and 30 Hz for offline filters (Range: 8.5 – 50 Hz). Eight studies (16.3%) used an online notch filter — all at 50 Hz— and three studies (6%) used an offline notch filter (50 Hz for two studies and 60 Hz for the other). For online referencing schemes, a single mastoid (left or right) was the most common choice, comprising 11 of the 49 included studies (22%), followed by linked ears (16.3%), Cz (12.2%) and the nose (10.2%). The online reference was not reported in 7 studies (14.28%). Unless the data were explicitly re-referenced, it was relatively uncommon for studies to report the offline reference separately, resulting in missing values for 30 studies (60%). The average reference was the second-most frequent choice, reflecting 12 studies (24%).

#### 4.1.5 ERP Reporting

Twenty-eight of the 49 included studies (57.14%) provided associations for both P3 amplitudes and latencies, while 13 studies (26.53%) reported associations only for amplitudes and eight (16.33%) only for latencies. Of studies that investigated amplitude associations, 20 (48.8%) noted any descriptive statistics for amplitude values, while 21 out of 36 studies (58.3%) on latency associations reported any descriptive statistics for latencies. Across electrodes and irrespective of the tasks during EEG recording (see next section), P3 amplitudes were numerically largest at Cz, followed by Pz, “other”, and Fz. P3 latencies peaked earliest for Fz, followed by Cz, “other”, and Pz (Table 3).

**Table 3.** Descriptive Statistics for P3 Amplitudes and Latencies by Electrode.

| *Variable* | *Electrode* | *Mean* | *SD* | *Min* | *Max* | *Studies* | *Samples* | *Effects* |
| --- | --- | --- | --- | --- | --- | --- | --- | --- |
| Amplitudes | Fz | 5.64 | 3.82 | -0.15 | 16.69 | 6 | 6 | 18 |
|  | Cz | 9.50 | 3.38 | 2.04 | 23.47 | 4 | 4 | 7 |
|  | Pz | 8.79 | 4.81 | 2.45 | 30.95 | 11 | 11 | 72 |
|  | Other | 8.22 | 3.13 | -0.45 | 12.40 | 9 | 9 | 75 |
| Latencies | Fz | 367.86 | 38.27 | 280.11 | 477.44 | 8 | 8 | 23 |
|  | Cz | 387.74 | 43.14 | 251.47 | 450.74 | 6 | 6 | 20 |
|  | Pz | 406.26 | 59.96 | 238.13 | 461.41 | 13 | 14 | 143 |
|  | Other | 396.00 | 43.10 | 265.00 | 485 | 6 | 6 | 37 |

*Note:* Mean, SD, min, and max, values reflect the mean values for each statistic across all available samples (e.g., min = average of the lowest reported amplitude across samples). Studies and samples reflect the number of studies and sub-samples reporting mean P3 values, as these were reported most frequently. Effects refers to the total number of mean values reported across all samples, experimental task conditions and electrodes in the current meta-analytic database. Other refers to any electrode(s) or electrode combination excluding Fz, Cz, Pz. Amplitudes are reported in microvolts. Latencies are reported in milliseconds.

Forty-one studies tested for potential associations between GCA and P3 amplitudes. Among these, 25 (61.0%) used peak amplitude, 11 (26.8%) reported mean amplitude measurements, and a single study used a cluster-based permutation test to operationalize amplitude effects (2.4%; Markiewicz et al., 2021). Finally, four studies (9.8%) did not clearly indicate their method of amplitude measurement. Thirty-six studies tested P3-GCA associations for latencies. Thirty-one of these examined peak latency measures (86.1%), with only a single study also mentioning fractional area latency measures (2.78%; Schubert et al., 2023). Five studies (13.89%) did not provide clear information about their method of measuring P3 latencies.

The most common interval for measuring the P3 (i.e., “What is the most common *time window*?”) was 250-550 milliseconds (ms) post-stimulus. This refers to 164 (16%) of amplitude associations and 164 (25.62%) latency associations. When measurement start and end times were considered separately, the most common measurement onset and offset times were 300 and 700 ms post-stimulus for amplitudes and latencies (12.8% and 19.2%, respectively) followed by 250 and 550 ms (7.7% for amplitudes, 11.5% for latencies). The earliest respective starting points for amplitude and latency measurements were 3 and 180 ms post-stimulus, and the latest ending point was in both cases 1000 ms. Overall, however, the included studies were characterized by substantial heterogeneity in measurement windows, with 34 and 23 unique start and end pairings for amplitude and latency measurements, respectively. Critically, seven studies measuring amplitudes (15.5%) and 16 studies measuring latencies (40%) did not clearly report their measurement windows

### 4.2 Study Quality Assessment (DIAD)

The global DIAD quality scores of each study are listed in Table 1 on the right side. More detailed results of the design and implementation questions, the contextual questions as well as the rating algorithm are provided in our online Supplementary Material: https://osf.io/j9g7q/.

#### 4.2.1 Construct Validity: Fit Between Concepts and Research Questions / Hypotheses

All 49 studies included in this meta-analysis demonstrated at least an adequate “fit between concepts and research questions / hypotheses” (Yes: 4, Maybe Yes: 45). This high consistency likely reflects the well-established operationalization of general cognitive ability (GCA) and the P300 ERP component within cognitive neuroscience. Detected flaws that led to a lower rating in this DIAD category (Maybe Yes instead of Yes) resulted in most cases from missing information on the reliability of P300 amplitude and latency measures (questions 1.4.3 and 1.4.4), which were often not reported in the original publications.

#### 4.2.2 Internal Validity: Clarity of Causal Inference

Ratings for “clarity of causal inference” of the included studies were mixed (Maybe Yes: 23, Maybe No: 25, No: 1). One study received an overall “No” rating because essential details concerning the EEG recording environment (question 2.1 and subquestions) were not reported. The majority of “Maybe No” ratings resulted from missing information regarding the administration of GCA assessments (question 2.2 and subquestions), for instance whether the test administrators were trained or whether the participants were fluent in the test language.

#### 4.2.3 External Validity: Generalizability of Findings

Ratings for “generalizability of findings” were overall moderate (Yes: 3, Maybe Yes: 28, Maybe No: 18). Although all studies included healthy adults, the range of participant characteristics and GCA measures was limited, leading to lower ratings in several cases.

#### 4.2.4 Statistical Validity: Precision of Outcome Estimation

The DIAD category statistical validity, precisely “precision of outcome estimation”, showed the largest variation and ratings were overall quite low (Maybe Yes: 6, Maybe No: 18, No: 15, not applicable: 10). Many studies received lower scores because statistical assumptions were not tested properly, or the selected measures of association were not appropriate to the data (questions 4.1.1 and 4.1.2). In addition, several studies did not report all test statistics appropriately, inclusively confidence intervals (questions 4.2.1 and 4.2.2). For ten studies, this whole category was rated as “not applicable” because the data on GCA and the P300 had to be obtained directly from the authors.

### 4.3 Meta-analytic Synthesis of Amplitude Effects

#### 4.3.1 Hypothesis 1: Positive Association between General Cognitive Ability and P3 Amplitudes

The across-study association between P3 amplitudes and GCA was small but significant: *r* = .13 (381 original study effect sizes from 46 samples, Fisher’s *z* = 0.13; SE = 0.033, *z* = 3.90, *p* < .0001, 95%-CI [0.064, 0.19]). Across-study heterogeneity was meaningful (Q(380) = 914.81, *p* < .0001) with I^2^ = 60.36% and between-sample variance of 0.042 (Figure 3).

**Figure 3.**
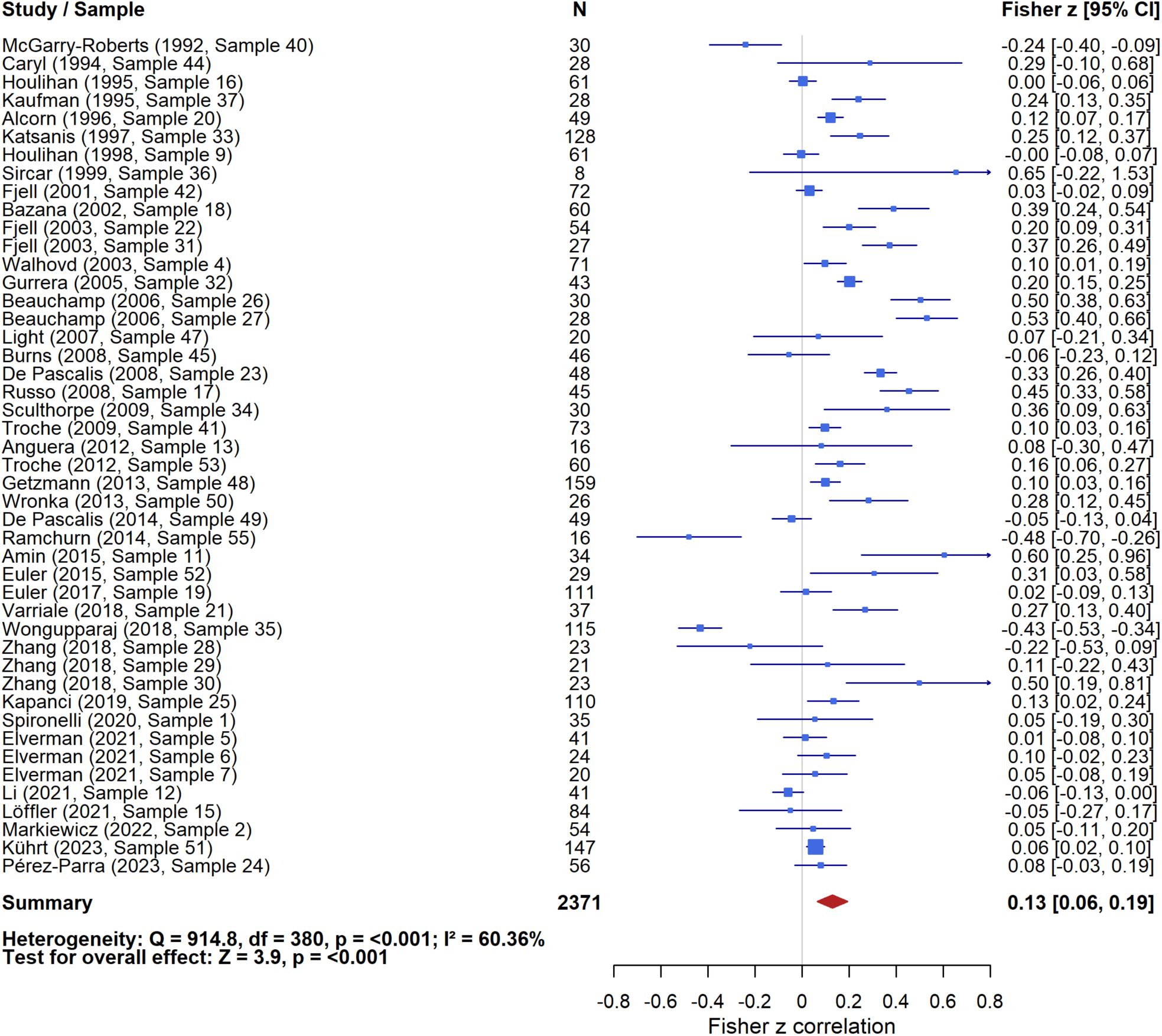
Significant but small positive across-study association between general cognitive ability and P3 amplitude. Forest plot of study sample-specific associations between individual differences in general cognitive ability (GCA) and variation in P3 amplitude. Each blue box represents the inverse-variance weighted mean Fisher’s *z* correlation for a given study sample; horizontal lines show 95% confidence intervals. One effect per sample is shown, computed by averaging all relevant sub-effects using inverse-variance weighting. Some studies contribute multiple samples. Box sizes reflect the relative influence of each sample on the overall meta-analytic estimate, based on that weight. The meta-analytically derived across-sample effect size is illustrated by a red diamond, and metrics of across-sample heterogeneity (Q, I^2^) together with statistics of the meta-analytic across-study effect (Fisher’s *z*) are presented at the bottom of the plot.

#### 4.3.2 Hypothesis 1A: Moderation Effect of Task Type

Concerning our hypothesis that task type moderates the association between P3 amplitude and GCA, such that the correlation is positive in oddball tasks and negative in tasks requiring greater levels of cognitive control, a significant moderation effect was observed (*QM*(4) = 17.75, *p* = .0014) with a significant positive association between P3 amplitudes and GCA in Oddball tasks (Fisher’s *z* = 0.26, SE = 0.043, *z* = 5.92, *p* < .001, CI [0.170 - 0.339]; based on 381 associations nested within 46 samples) and significantly smaller associations for Attentional/Cognitive Control tasks (Δ*z* = –0.24, SE = 0.062, *z* = -3.90, *p* < .0001), Working Memory tasks (Δ*z* = –0.20, SE = 0.064, *z* =-3.17, *p* = .0015), Chronometric tasks (Δ*z* = –0.25, SE = 0.090, *z* =-2.81, *p* = .0049), and “Other” tasks (Δz = –0.19, SE = 0.076, *z* = -2.56, *p* = .0104). However, in contrast to our hypothesis, effects for tasks requiring cognitive control were, although significantly smaller and very close to zero, also positive (Figure 4). Significant residual heterogeneity remained after accounting for variance due to Task Type *QE*(376) = 775.07, *p* < .0001, with between-sample variance σ² = 0.030 and an associated *I²* = 53%.

**Figure 4.**
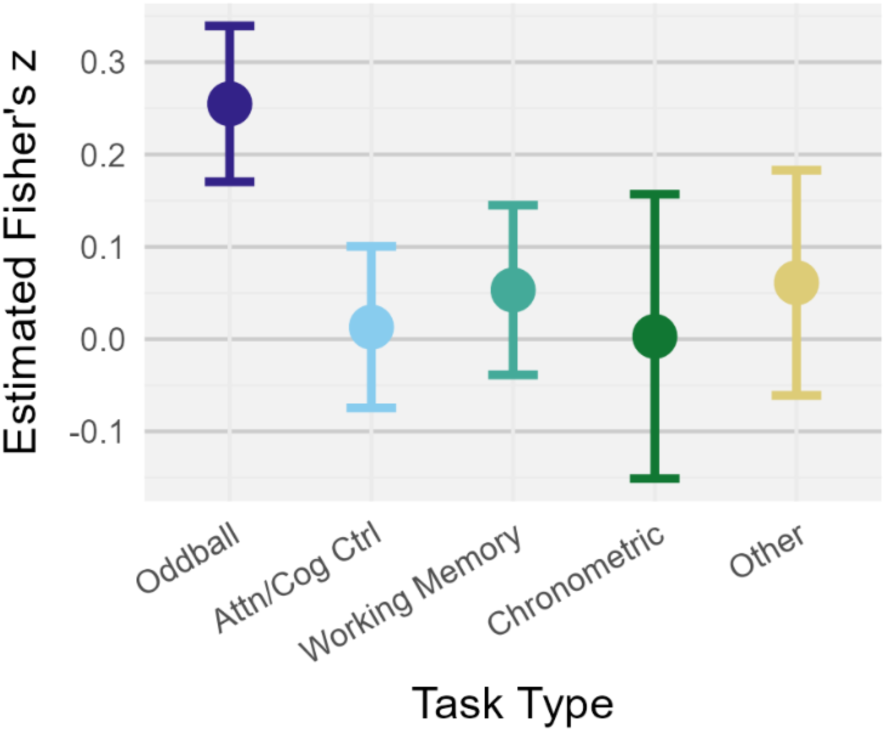
Task type moderates the across-study associations between general cognitive ability and P3 amplitude. Error bars reflect 95% confidence intervals on marginal means.

#### 4.3.3 Hypothesis 1B: Moderation Effect of Task Difficulty

With respect to our hypothesis that task difficulty moderates the association between GCA and P3 amplitudes, no significant effect was observed (*QM*(2) = 3.45, *p* = 0.18; 141 associations from 16 samples). The estimated Fisher’s *z* for the across-study P3-GCA association for low difficulty tasks was 0.153 (SE = 0.054, z = 2.86, *p* = 0.004), with non-significant increases for medium difficulty tasks (Δ*z* = 0.050, SE = 0.027, z = 1.85, *p* = .065) and high difficulty tasks (Δ*z* = 0.019, SE = 0.034, z = 0.578, *p* = .563). The between-sample variance was σ² = 0.038, with an associated *I^2^* = 63.9%.

Exploratively, we also examined whether the mean amplitude of the P3 reported in all included studies differed between low, medium and high difficulty tasks, irrespective of its association with GCA. These values were reported relatively infrequently, yielding 74 observations drawn from 7 samples. The estimated P3 means in microvolts (±95% CI) were: 6.23 [3.72, 8.74] for low difficulty tasks, 6.25 [3.74, 8.76] for medium difficulty tasks, and 6.25 [3.74, 8.76] for high difficulty tasks. Pairwise differences were small and non-significant, with substantial residual heterogeneity *QE*(71) = 13,832, *p* < .0001.

#### 4.3.4. Examination of Outliers

A single study (Wongupparaj, et al., 2018) was identified as potential outlier with respect to GCA-P3 amplitude associations based on the joint criterion of studentized residuals ± 1.96 (Viechtbauer & Cheung, 2010), and a Cook’s distance value larger than the median value plus six times the interquartile range (0.145 for the global amplitude model). We confirmed that the study met our inclusion criteria and the data were extracted properly. As this was the case, we refrained from excluding it *a priori* but performed a post-hoc sensitivity analysis of the global amplitude model excluding this study. The direction and significance of the main effect was unchanged, although the magnitude increased (to *r* = .14 from *r* = .13) and between-sample variance (σ²) decreased from .042 to .033.

#### 4.3.5 Publication Bias

Tests for publication bias provided no consistent evidence of funnel plot asymmetry (Figure 5). Egger’s regression test was not significant (*z* = 1.29, *p* = 0.20), nor was Begg’s rank correlation test (*τ* = 0.13, *p* = 0.22). Trim-and-fill imputation suggested up to two studies potentially missing on the left side of the funnel (inverse-variance weighted SE ≤ 0.45). After imputing these studies, the adjusted estimate of the meta-analytical P3-GCA amplitude association across all studies was virtually unchanged and remained significant (*z* = 3.59, *p* = 0.003, 95% CI [0.054, 0.183]). A fail-safe *N* analysis indicated that 114 unpublished null studies would be required to reduce the overall effect to non-significance.

**Figure 5.**
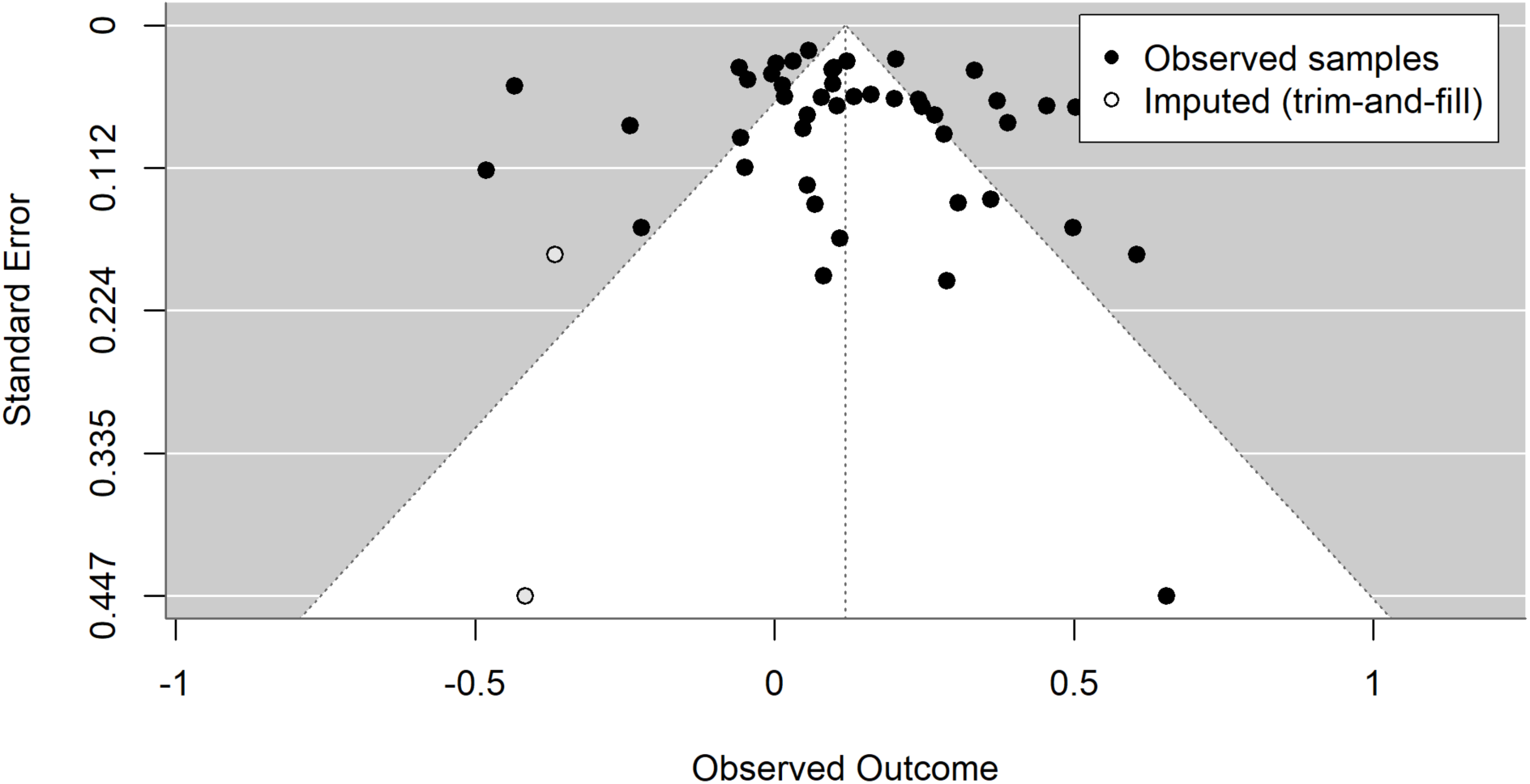
No support for potential publication bias in the across-study association between general cognitive ability and P3 amplitude. Funnel plot of P3–GCA amplitude associations (Fisher’s *z*) vs. standard error with trim-and-fill. Each point represents one study sample; multiple effects within a sample were combined using a fixed-effect inverse-variance weighted mean on the Fisher’s *z* scale. Trim-and-fill imputed two studies (open markers).

### 4.4 Meta-Analytic Synthesis of Latency Effects

#### 4.4.1 Hypothesis 2: Negative Association between General Cognitive Ability and P300 Latency

Analyses of our second primary hypothesis of a negative association between GCA and P3 latency resulted in a meta-analytic across-study effect size of -0.184 (SE = 0.030, *z* = -6.231, *p* < .0001, 95% CI [-.242, -0.126]; 336 associations, nested within 40 samples). This corresponds to a pooled correlation of *r* = -0.18, thus, supporting our hypothesis. The estimated between-sample variance σ² was 0.028, indicating meaningful heterogeneity across samples (*Q*(335) = 852.15, *p* < .0001), with an associated *I^2^* = 52.20% (Figure 6).

**Figure 6.**
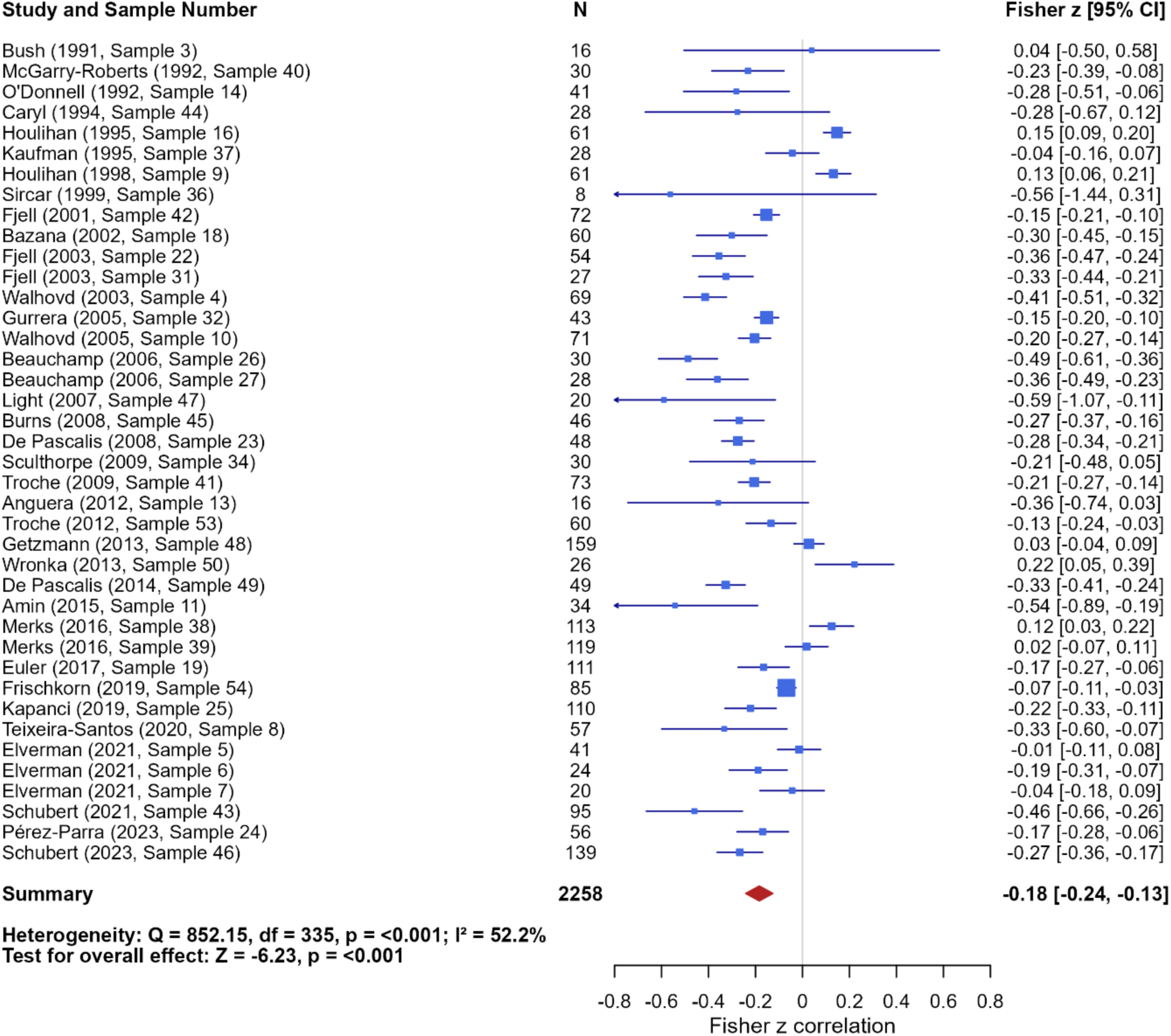
Significant but small negative across-study association between general cognitive ability and P3 latency. Forest plot of sample-specific associations between individual differences in general cognitive ability (GCA) and variation in P3 latency. Each blue box represents the inverse-variance weighted mean Fisher’s *z* correlation for a given study sample; horizontal lines show 95% confidence intervals. One effect per sample is shown, computed by averaging all relevant sub-effects using inverse-variance weighting. Some studies contribute multiple samples. Box sizes reflect the relative influence of each sample on the overall meta-analytic estimate, based on that weight. The meta-analytically derived across-sample effect size is illustrated by a red diamond, and metrics of across-sample heterogeneity (Q, *I^2^*) together with statistics of the meta-analytic across-study effect (Fisher’s *z*) are presented at the bottom of the plot.

#### 4.4.2 Hypothesis 2A: Effect of Task Difficulty

Our final *a priori* hypothesis that the magnitude of the association between GCA and P3 latency would increase with increasing task difficulty was not supported by our analyses (*QM*(2) = 3.31, *p* = 0.19; 151 associations from 17 samples). The association between GCA and P3 latency was significantly negative for tasks of low difficulty (Fisher’s *z* estimate = -0.12, SE = 0.052, *z*-value = -2.279, *p* = 0.023, 95% CI [–0.220, -0.017]), while it was non-significantly negative for medium difficulty tasks (Δz = –0.054, SE = 0.031, *z*-value = -1.722, *p* = 0.085) and high difficulty tasks (Δz = -0.047, SE = 0.035, *z*-value = -1.325, *p* = 0.185). These difficulty levels did not differ significantly from each other. Substantial residual heterogeneity remained, *QE*(148) = 464.44, *p* < .0001, with between-study variance estimated at σ² = 0.027, and approximate residual *I^2^*= 58.7%.

Finally, as we did for P3 amplitude, we exploratively examined whether task difficulty was related to mean P3 latencies, irrespective of its association with GCA. Latencies were significantly longer in medium difficulty tasks (Δ = 50.27 ms, *p* < .0001) and high difficulty tasks (Δ = 44.32 ms, *p* < .0001), relative to low-difficulty tasks (M = 397.95 ms, SE = 23.02, *z* = 17.29, *p* < .0001, 95% CI [352.83, 443.06]; 107 associations nested within 10 samples; and see Figure 7). The omnibus test of mean differences by Difficulty was significant, QM(3) = 1721.68, *p* < .0001. Substantial heterogeneity remained QE(104) = 13,114.10, *p* < 0.001, with variance components σ² for between-sample heterogeneity = 5094.71, and σ² for electrode-level heterogeneity nested within sample = 205.83.

**Figure 7.**
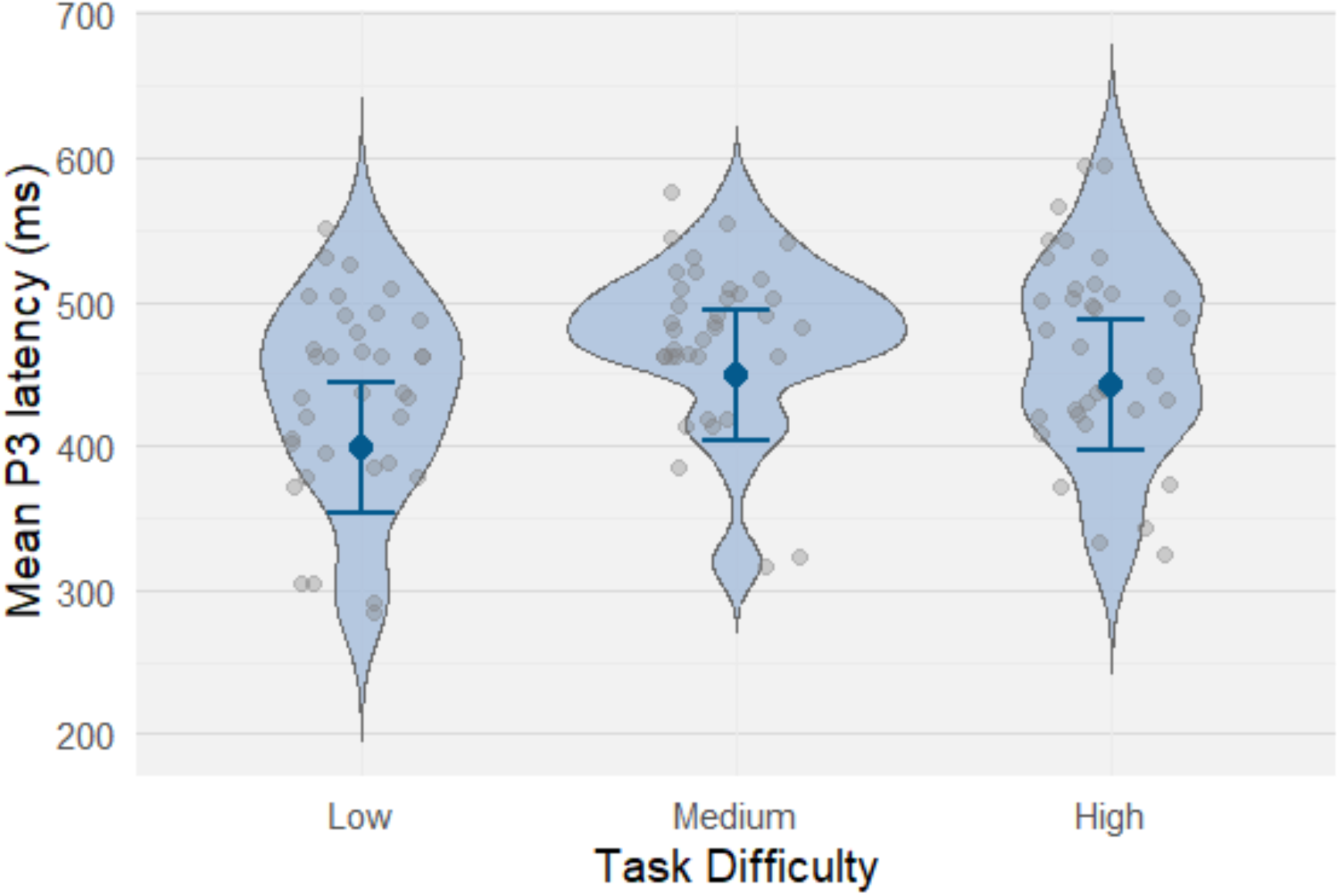
Mean P3 latencies as a function of difficulty, irrespective of their association with GCA. Blue dots and bars reflect model-estimated means and 95% confidence intervals. Gray dots reflect individual sample values.

#### 4.4.3 Examination of Outliers

No study was identified as potentially influential based on the joint criterion of studentized residual greater than ±1.96 and Cook’s distance value greater median plus six times the interquartile range (Viechtbauer & Cheung, 2010), which was 0.18 for the global meta-analytical association between GCA and P3 latency. Using a less conservative cut-off of a Cook’s distance of > 4/k (0.10 for this model with k = 40 clusters), two studies were considered influential (Houlihan et al., 1994; Beauchamp & Stelmack, 2006). When the model was re-run without these studies, the direction and significance of the main effect were unchanged, the pooled estimate was virtually identical, and the between-sample variance (σ²) decreased modestly from .028 to .023.

#### 4.4.4 Examination of Publication Bias

Tests for publication bias on the meta-analytical across-study association between GCA and P3 latency suggested evidence of funnel plot asymmetry (Figure 8). Egger’s regression test was significant (*z* = –2.31, *p* = .021), while Begg’s rank correlation test was not significant (*τ* = – 0.10, *p* = .38). Trim-and-fill imputation indicated eight studies potentially missing on the right side of the funnel (largest observed inverse-variance weighted SE = 0.45). After imputing these studies, the adjusted meta-analytical association between GCA and P3 latency remained significant (Fisher’s *z* = –0.138, *p* < .001, 95% CI [–0.198, –0.078]). A fail-safe *N* analysis indicated that 288 unpublished studies with null effects would be required to reduce the overall effect to non-significance. Influence analyses showed that no single study unduly influenced the pooled estimate or heterogeneity; the most imprecise study (n = 8) did not account for the observed funnel plot asymmetry, which remained significant after excluding it.

**Figure 8:**
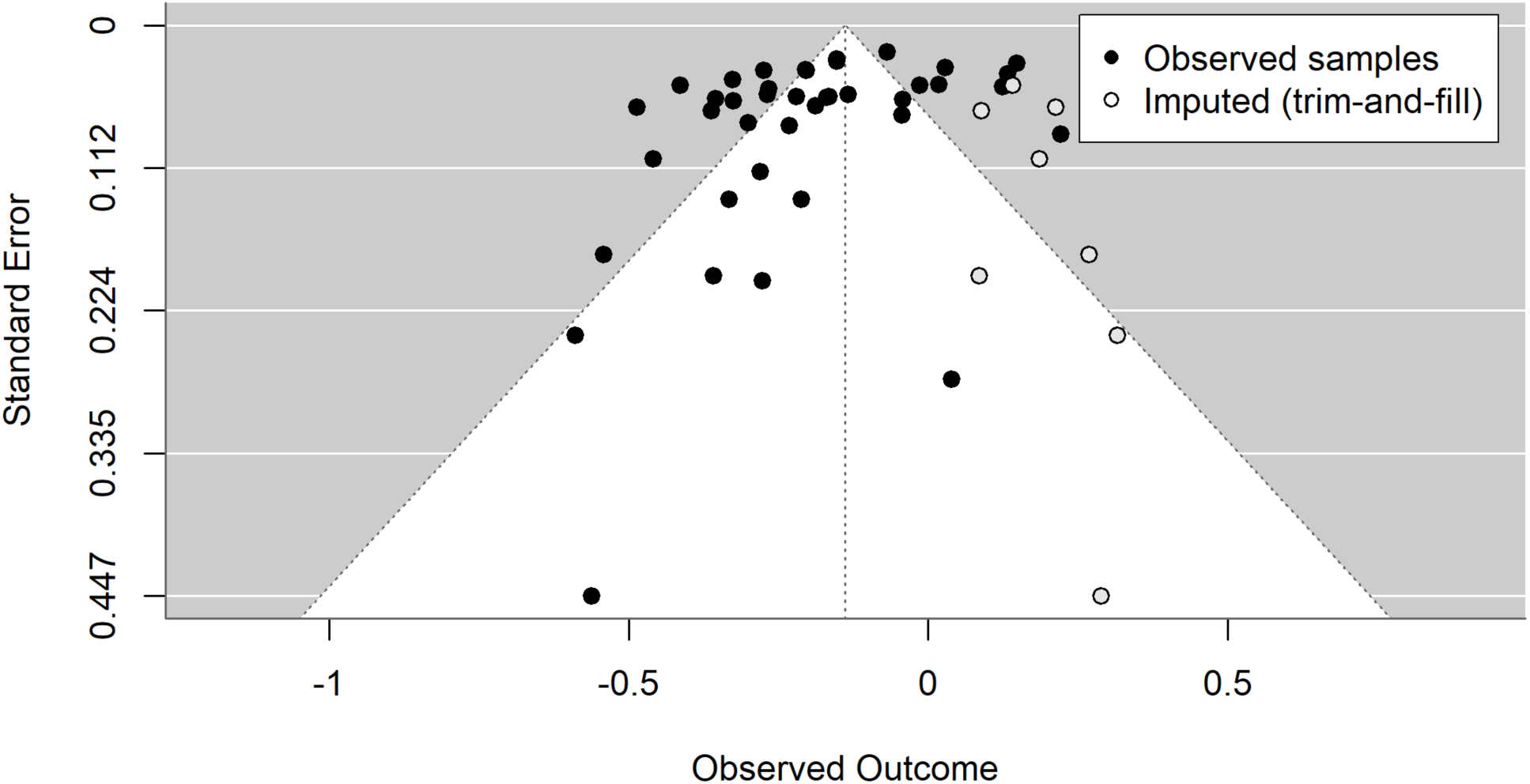
Support for presence of publication bias in the across-study association between GCA and P3 latency. Funnel plot of P3–GCA latency associations (Fisher’s *z*) vs. standard error with trim-and-fill. Each point represents one study sample; multiple effects within a sample were combined using a fixed-effect inverse-variance weighted mean on the Fisher’s *z* scale. Trim-and-fill imputed seven studies (open markers).

## 5. Discussion

This preregistered study presents the first systematic review evaluating the state and quality of the literature on the relationship between the P3 event-related brain potential (ERP) and general cognitive ability (GCA). Furthermore, it presents the first quantitative meta-analysis determining whether P3 amplitudes and latencies are significantly associated with individual differences in GCA. We identified 49 studies that contributed up to 381 effects (out of 5641 screened studies), which were eligible for PRISMA-based meta-analytic comparison and rated study quality with the Study Design and Implementation Assessment Device for Individual Difference Research (DIAD-ID). In accordance with our hypotheses, a small but significant positive across-study association was observed for general cognitive ability and P3 amplitudes (*r* = .13; 95%-CI [.06, .19]), while a significant negative across-study association was seen for P3 latencies (*r* = -.18; 95%-CI [-.24, -.13]). Heterogeneity between studies was substantial, sub-analyses highlighted potential moderators such as type of the task during EEG recording, and we identified potential publication bias concerning studies on the association between GCA and P3 latencies. In the following, we highlight limitations, open questions, and provide guidelines to increase the comparability in future studies on the neurobiological underpinnings of individual differences in cognitive ability.

### Quality of Existing Empirical Evidence

The systematic review indicated that most samples were comprised of young, right-handed, relatively well-educated, and predominantly female participants. For studies reporting IQ scores, sample participants tended to score near the high-average range, with less variability within samples compared to population norms. Most studies used measures of fluid or general intelligence (∼61%), which reflects the focus on these constructs compared to, e.g., crystallized intelligence, in the field. Nearly two-thirds of studies (∼64%) elicited the P3 via oddball or cognitive control tasks, reflecting the P3’s history as indicator of higher-level cognitive processes (Luck, 2012). There was substantial heterogeneity in the literature concerning recording, signal processing, and measurement approaches, which may have contributed to inconsistent findings and challenges replicability (Clayson et al., 2021; Soskic et al., 2022). This “Garden of Forking Paths” (Gelman & Loken, 2014) presents a well-known problem in EEG research, but needs to be overcome to increase the reliability and validity of findings. Common reporting standards, greater transparency, and standardization are urgently required (Trübutschek et al., 2024). Most commonly, the reviewed studies used peak amplitude and latency measures to examine associations with GCA from 250-550 ms post-stimulus at electrode Pz. While the respective window and the electrode Pz present reasonable choices to reliably measure the P3 (Luck, 2012, 2014), peak amplitude and latency are characterized by important limitations such as susceptibility to high-frequency noise. Thus, while it can be valuable to report peak amplitude and latency for comparability with previous research, mean amplitude and fractional area latency have been shown to reflect preferable candidates as primary outcome measures (Luck, 2014).

To systematically assess study design quality, we developed the Study Design and Implementation Assessment Device for Individual Difference research (DIAD-ID) based on the DIAD from Valentine & Cooper (2008). Our system included four broad outcome category questions that inform about construct validity, internal validity, external validity and statistical validity, 11 composite questions, and 36 more specific design and implementation questions as well as a specific algorithm detailing how to synthesize lower-level ratings into higher level outcomes. First, studies were rated relatively highly with respect to construct validity, such as the operationalization of GCA and the P3. This was likely driven by the fact that we per-se focused our review on studies applying established GCA measures and studies providing clear P3 measures, thus automatically excluding studies which very low construct validity. Indeed, during screening, 44 studies were excluded due to the use of non-established GCA measures, four studies due to unstandardized measures, and one due to a self-reported measure of GCA, thus suggesting much lower construct validity in the whole literature published on GCA-P3 associations than can be inferred from this observation.

Second, study quality ratings were mixed on the dimension of internal validity—typically due to missing information about aspects of EEG recording or GCA measurement procedures, again reflecting the need to adhere to reporting standards and increase transparency (see above). Third, ratings were also mixed for external validity and generalizability, largely owing to limited variability in sample demographics (primarily female young healthy college students) and GCA measures (global and fluid intelligence). Finally, the category of statistical validity was rated most variably and typically lowest among the dimensions considered. This was primarily driven by lacking tests of statistical assumptions, inappropriateness of measures, and missing reports of statistics—inclusively confidence intervals.

### Higher GCA is Associated with Higher P3 Amplitudes

A total of 46 samples (381 correlations) contributed to the global meta-analysis of associations between P3 amplitudes and GCA. Consistent with our preregistered hypotheses, the results confirm a small but significant positive association between GCA and amplitudes. This suggests that in general, higher cognitive ability is associated with greater neural activation as indexed by the P3. However, as also negative associations have been reported in some studies, the finding of a small overall effect was not unexpected. Nevertheless, the presently reported effect size of *r* = .13 represents an important benchmark for future studies to further explore and expand upon.

### Functional Significance of the GCA-P3 Amplitude Association

The amplitude of an ERP component depends on the both the organization of the involved neural generators and the degree to which their activity is synchronized in response to an eliciting event (Fabiani et al., 2012; Luck, 2014). Accordingly, variability in component amplitudes across tasks or individuals reflects differences in the extent and configuration of this event-related neural activity. The key question raised by the present findings, then, is why such variability should relate to GCA at all, and why that relationship might be positive.

Three of the most prominent theories on the functional significance of the P3 variously hold that it represents (a) the neurocognitive revision or “updating” of the current representation of the state of the environment (Donchin, 1981) or information held in working memory (Luck, 2014), (b) the end of a process of sensory evidence accumulation that underlies decision-making (Twomey et al., 2015), or (c) the reactivation of previously-established stimulus-response links (S-R link hypothesis; Verleger, 2020). Clearly, all these theories share features with important findings or accounts of GCA. For example, GCA has been conceptually and empirically linked to the ability to implement effective mental models of the world (Duncan, 2010; Duncan 2025), to individual differences in working memory capacity (Colom et al., 2004; Kyllonen, & Christal, 1990; Oberauer et al., 2005; Süß et al., 2002), and to the efficacy of, and constraints set by, perceptual processing (Acton & Schroeder, 2001; Galton 1883; Grudnick, & Kranzler, 2001; Melnick et al., 2013). A potential connection between GCA and the S-R link hypothesis can also be established: In brief, the S-R link hypothesis holds that the P3 reflects the reactivation of an already-established but dormant stimulus-response mapping; for example, between the infrequent target stimulus and the previously-established correct response in an oddball task. This differs from the context-updating account in proposing a more specific process underlying the P3, and by emphasizing an existing stimulus-response mapping rather than a revision. That said, insofar as the S-R link hypothesis implicitly involves elements like the need to keep track of environmental contingencies, and the need to expend neural resources to do so, the S-R link hypothesis may connect the P3 to accounts of GCA that involve maintaining mental models of the world (e.g., Duncan et al., 2025).

From the perspective of theories focused on individual differences in GCA or intelligence, at least two additional concepts offer plausible connections to P3 amplitudes: The first is connectivity, where both structural and functional connectivity in fronto-parietal and other distributed networks have long been emphasized as a potential basis for individual differences in GCA (Jung &, Haier, 2007; Duncan, 2010; Basten et al., 2015; Barbey, 2018; Thiele et al., 2025; for review: Hilger & Sporns, 2021), and which likely overlap with the distributed neural generators of the P3 (Bocquillon et al., 2011; Linden, 2005). In addition, various microscale features, like dendritic arborization and neurite density (Genç et al., 2018; Goriounova et al., 2018), have also been identified as correlates of GCA and may influence larger-scale networks associated with both GCA and the P3. Therefore, we propose that enhanced connectivity among relevant networks at various scales might enable greater synchronization in the neural processes governing event-related mental updating, sensory evidence accumulation, or re-activation of stimulus-response contingencies, and may thus contribute to the P3 amplitude associated with GCA.

A second concept that may inform the GCA-P3 amplitude link is predictive processing. In brief, the account of predictive processing holds that the brain continuously tries to predict both the organism’s internal states and the external state of the world. The goal is to minimize deviations from expected states. Only the latter, the deviations, are processed (costing energy and time) and give rise to neuronal activity to update the internal model (i.e., prediction error minimization; Friston, 2010; Clark, 2015). Of note, the P3 was initially proposed as reflecting sensory prediction errors (Friston, 2005), which resonates with both the context updating and S-R link hypotheses of the P3. However, research directly supporting this assumption is lacking so far (see Verleger, 2020, pp. 5-7). This may potentially be due to different manifestations of the P3 relating to very specific aspects of the error minimization process (Barceló, 2021; Kopp, 2008; Kopp et al., 2016; Wacongne et al., 2011) – a possibility that should be explored in future research.

Predictive processing has also been proposed as critical to individual differences in GCA and intelligence (Euler, 2018; Trapp et al. 2021, 2025). Euler (2018) proposed that the proposed neural hierarchy of lower- to higher-order predictions may, in part, map on to the narrow-to-broad cognitive hierarchy observed in psychometric research on intelligence (Carroll, 1993; Johnson & Bouchard, 2005) and that greater P3 amplitudes may relate to higher GCA because they reflect faster or more coordinated activity to carry out a specific cognitive process related to both GCA and predictive processing (e.g., belief updating; Barceló, 2021; Kopp et al., 2016). Trapp and colleagues (2025) argued further that intelligence not only requires accurately predicting sensory states, but also correctly estimating the *precision* of those predictions. This would inform the perceptual decision-making account of the P3, but again direct empirical evidence is missing. Overall, we propose these frameworks as promising bases for future research on the association between P3 amplitudes and GCA.

### The GCA-P3 Amplitude Association is Moderated by Task-Type but not by Task Difficulty

Beyond the existence of a positive association between GCA and the P3, we hypothesized that this association is moderated by task type such that oddball tasks produce positive associations, and more mentally demanding tasks result in negative associations. While a significant moderation effect was present with oddball tasks eliciting significant positive associations with GCA, more demanding tasks resulted in much smaller associations close to zero, rather than reliably negative. This is contrary to our hypothesis and seems to contrast with substantial previous work emphasizing the importance of more demanding cognitive processes for intelligence (Binet & Henri, 1895; cited in Siegler, 1992; Conway et al., 2003; Gottfredson, 1997; Kovacs & Conway, 2016). It also contrasts with very recent proposals stating that the use of trait-relevant tasks (here more demanding tasks) during neuroimaging increases the detectability of associations between a given trait and its neural correlates (here GCA; Popp et al., 2025a, b, 2026; DeYoung et al., 2025; Thiele et al., 2025). One explanation for our unexpected finding could lie in the fact that the oddball tasks included in our meta-analysis were rather homogeneous, while more demanding tasks differed considerably among each other. In turn, this could have produced greater variability in the processes elicited by each task and increased the variability in how the P3 precisely manifests (including interactions and distortions by other processes; e.g., Kok, 2001), making it ultimately more difficult to identify a significant across-study association. Indeed, recent work has convincingly shown task specificity of the GCA-P3 relationship for latencies (see below), which may apply to amplitudes as well (Sadus et al., 2025) and requires further research.

Irrespective of its explanation, the observed moderation has specific implications. First, it shows that it is important to explicitly consider the type of the task when comparing P3-GCA effects across studies to prevent over- or underestimation of the “true” effect size. This not only concerns studies on GCA but studies on neural correlates of individual differences in general (DeYoung et al., 2025) – tasks need to be matched to the trait of interest thoroughly. Second, the fact that the magnitude, if not the direction, of the relationship varies by task may also help explain why associations for amplitudes have been more inconsistent than effects for latencies in the GCA literature (Euler & Schubert, 2021).

Finally, our second moderation hypothesis, that the association between P3 amplitude and GCA would be greater in more difficult tasks, was not supported by our analyses. Several factors could have contributed to that null finding. First, only a comparatively small number of studies provided data to address this question (*n* = 17 samples). Second, these samples varied considerably in terms of the tasks they used to elicit the P3, and in turn, the levels of difficulty they manipulated. That this may have contributed to the observed null-finding is supported by our explorative post-hoc analysis in which could not replicate the previously reported general effect of difficulty on mean P3 amplitudes in our sample (irrespective of GCA differences; Kok, 2001). Third, although difficulty has been shown to affect *mean* P3 amplitudes by previous work, which represent within-subjects effects, this may not bear on the between-subjects relationship of the P3 to GCA. We therefore conclude that the systematic identification of task attributes systematically shaping GCA-P3 amplitude association presents an interesting subject for further study. Finally, there was little evidence that these findings were significantly influenced by publication bias.

### Higher GCA is Significantly Associated with Faster P3 Latencies

A total of 40 samples (336 correlations) contributed to the global meta-analysis of associations between P3 latencies and GCA. Consistent with our hypotheses, we detected a small but significant negative relation, suggesting that in general, higher cognitive ability is associated with faster neural responses to specific stimuli. However, similar but not as frequently as for the association between GCA and P3 amplitudes, some studies reported also associations of opposite direction, i.e., longer latencies for higher GCA. Thus, we consider the resulting effect size of *r* = -.18 as a meaningful benchmark for future research.

### Functional Significance of the GCA-P3 Latency Association

As the first systematic review and meta-analysis of the GCA-P3 latency relation, the current findings provide important support for the mental speed hypothesis of intelligence (Jensen, 2006). In brief, this hypothesis holds that individual differences in the speed of mental operations are pre-eminent among lower-order constructs in generating individual differences in GCA, relative to alternative constructs like relational processing (Chuderski, 2022; Jastrzębski, et al., 2020), working memory, and other executive abilities (Kovacs & Conway, 2016). Support for this hypothesis comes from studies using “chronometric” tasks like reaction time and inspection time (Deary et al., 2001; Sheppard & Vernon, 2008; Troche & Rammsayer, 2009), and studies demonstrating moderate to large associations between GCA and ERP latencies in latent factor models (Schubert et al., 2015; Schubert et al., 2017; Schubert et al., 2023).

At the neural level, greater mental speed has been related to the integrity of white matter tracts (Penke et al., 2012; Kuznetsova et al., 2016), structural features of specific regions involved in decision making (locus coeruleus and dorsal cingulate; Eckert, et al., 2023), and variation in neural oscillatory frequencies (Ociepka et al., 2022). Regarding the P3, the relation between faster latencies and higher GCA has been hypothesized to reflect greater efficiency of information transmission among frontal and temporo-parietal networks (Schubert et al., 2017). Notably, although we did not hypothesize that the negative relation between GCA and P3 latencies would be moderated by task type (and thus did not examine this), recent evidence suggests that the latency of the P3 in *decision making tasks specifically* is strongly related to GCA, but potentially not during other processes like working memory encoding (Sadus et al., 2025). Therefore, additional research on factors like task type and specific cognitive operations is essential. Interestingly, the latter findings regarding the decision-related P3, along with those relating neural processes of decision-making to mental speed (Eckert et al., 2023), may inform about the evidence accumulation account of the P3 (Twomey et al., 2015).

### The GCA-P3 Latency Association is Not Moderated by Task Difficulty

Based on rather substantial literature (e.g., Kok, 2001; Polich, 2012), we hypothesized stronger GCA-P3 latency associations in more difficult tasks. However, our meta-analytical findings did not reveal a significant moderation. The same factors as discussed in respect to amplitude (see above) may have contributed to this null finding. However, unlike the analysis for amplitudes, our explorative post-hoc analysis identified a significant association between task difficulty and mean P3 latencies (irrespective of individual differences in GCA), with medium and high difficulty task conditions having longer latencies on average compared to low difficulty conditions, again highlighting the need for further research on moderating factors.

Finally, in respect to studies reporting GCA-P3 latency associations, there was some evidence of publication bias, and trim-and-fill analyses suggested that seven studies may be missing from the right side of the funnel plot. Nevertheless, the primary association remained significant after imputing these studies and a fail-safe *N* analysis suggested 288 unpublished null results would be needed to reduce the overall effect to non-significance.

## Limitations

Our meta-analysis has limitations. First, included studies were methodologically heterogeneous, and many had obtained the GCA-P3 associations only secondarily relative to their primary aims. Thus, relatively few studies were available to address our moderation hypotheses and more-specific exploratory moderation hypotheses would have been underpowered. Compelling subjects for future work include effects of different sub-domains of GCA (e.g., fluid versus crystallized intelligence or processing speed), examining associations between GCA and the P3 as recorded from standard versus target conditions in the oddball task, or from the target versus novelty P3, and examining interactions of specific reference schemes or other EEG-related features. To facilitate such future research, we provide our dataset online (https://osf.io/j9g7q/overview). This can easily be used as foundation for future meta-analyses that integrate additional data. Third, the observed across-study associations are only of small effect sizes, limiting the amount of variance in GCA that can be explained by the P3. In this regard, it is important to emphasize here that the upper bound of a detectable association is naturally limited by the (un)reliabilities of the behavioral and neural measures. Thus, since all measures have less than perfect reliability, obtained correlations will always underestimate the size of any true relationships. Latent variable modelling has been proposed as a solution to this issue (Schubert et al., 2023) but has not yet been widely adopted. Fourth, like the majority of research on neural correlates of GCA, our findings appear to imply differences in “lower-level” (rather than cognitive) processes between higher and lower-ability individuals. Contrastingly, it is also possible that the observed effects primarily or at least to some extent reflect differences in how individuals *approach* experimental tasks (e.g., differences in task engagement or strategies), and should be explored in future studies.

## Recommendations and Standards for Future Studies

The current review suggests recommendations for future work. First, in many cases it was difficult to identify even key parameters of the GCA-P3 associations (e.g., sample size, electrode of measurement), thus it is essential that future studies significantly improve their reporting praxis. Second, although sample sizes have improved over time, small and unrepresentative convenience samples dominated the literature reviewed here. The field would benefit greatly from increased efforts to recruit large and representative community samples and from sharing this data online. In this respect, the present results also provide a basis for formal power analyses to justify study sample sizes prior to recruitment. Third, only few studies reported reliabilities of their measures, or assessed GCA (or sub-domains) via multiple measures. However, as noted in the limitation section, knowledge about reliabilities (and use of multiple measures) is essential to (a) define the upper-bound of identifiable associations, and (b) distinguish between associations at the level of specific *tests*—which include both idiosyncratic and error variance—versus well-operationalized psychological *processes* (Colom & Thompson, 2011). Fourth, having established that not all versions of the P3 are equivalent for purposes of individual differences research, additional work is needed to understand how moderators like task type alter relationships with GCA. Specifically, single measurements of the P3 typically show both low reliability and high task-specificity (Schubert et al., 2023). This can be addressed by using multiple tasks to elicit the P3 and by then extracting latent variables capturing their shared variance (Euler & Schubert, 2021), while also matching tasks on the key cognitive operations of interest to assess potential moderation.

To increase progress in the field better comparability between studies is crucial. Future studies should therefore aim to converge on a common set of measurement parameters and on tasks or cognitive operations during which P3-GCA associations are measured. To this aim we propose the following concrete standards: Wherever experimental requirements permit, we suggest using (a) a measurement window of at least 250-550 ms post-stimulus (extending to 700 ms where possible), (b) a minimal montage of electrodes Fz, Cz, and Pz to assess the P3 relative to (c) an average reference, for maximal consistency with the existing literature. Furthermore, researchers should report (d) descriptive statistics on the sample (at least age, sex, handedness, education) as well as (e) descriptive statistics (at least mean, median, standard deviation, skewness) of both P3 amplitude and latency measures. As amplitude measures, we recommend using (f) mean and peak amplitude, the former because of higher validity (Luck, 2014), the latter to keep comparability with previous research. In respect to latency, the report of (g) fractional area latency (50%) and peak latency is advisable for the same reasons. Concerning the GCA-P3 association, future studies should (h) include scatterplots at least as supplements, and (i) use appropriate test statistics after checking all statistical assumptions thoroughly.

Finally, there exists some tension between the need to employ idiosyncratic tasks to address unique research questions and the need for greater comparability, clarity, and consistency in study methodology. Acknowledging that, we recommend (j) the development of one or two standard tasks that should be implemented (in additional to study-specific idiosyncratic tasks) in any future research on GCA-P3 associations. Valuable candidates for these two standard tasks might be a decision-making task in oddball and non-oddball variants (with the former also eliciting the novelty P3), or a task involving memory encoding (Sadus et al., 2025). Sharing these tasks, together with standard processing pipelines via established task libraries (e.g., ERP CORE; Kappenman, et al., 2021) or OSF, presents a promising means to enhance across-study comparability of GCA-P3 research to ultimately foster our understanding of general cognitive ability.

## Summary and Conclusions

This study provides the first systematic review and meta-analysis on the association between the P3 and general cognitive ability. Previous studies were characterized by convenience samples and heterogeneous methodology, reflecting a mix of studies addressing the GCA-P3 association primarily and studies reporting GCA-P3 relevant data secondarily to other aims. Studies varied in study design quality, with relatively high construct and external validity and low internal and statistical validity, while methodology improved over time. Meta-analytically, our results indicate a small, but significant, positive association between GCA and P3 amplitudes, as well as a small, but significant, negative association between GCA and P3 latencies. There was little evidence that the observed associations were meaningfully distorted by publication bias. For amplitudes, the association with GCA was moderated by task type, and strongest in the oddball task, indicating that eliciting conditions systematically influence observed relations. In contrast, task difficulty did not moderate the association with either amplitudes or latencies, though cross-study heterogeneity limited the strength of these analyses. Together, these findings suggest that relationships between the P3 and general cognitive ability exist but are neither large nor uniform. Future progress in this area will thus depend on increasing reporting praxis, the development and the use of common standard tasks, the recruitment of larger and more representative samples and finally, the implementation of open science practices. In providing numerical bases for power-calculations, in proposing concrete standards, and in sharing all data, analyses code, and a study quality rating scheme via OSF, this meta-analysis provides an important step along these paths to ultimately come closer to understanding the brain basis of intelligence.

## Author contributions

**Matt Euler:** Conceptualization, Data curation, Formal analysis, Funding acquisition, Methodology, Project administration, Writing - Original Draft, Visualization. **Kirsten Hilger:** Conceptualization, Data curation, Funding acquisition, Methodology, Project administration, Supervision, Writing – Original Draft.

## Acknowledgements

Our sincere appreciation goes to all authors that provided data as well as to Luisa Grittner and particularly Tobias Nöth for their great help in this project.

