## Supplementary figures and images for "Human Cognitive Ability and the P300 Event-Related Brain Potential: A Systematic Review and Meta-Analysis" (from another version of this preprint: version 1, 2026-02-16)

### Fig. 1

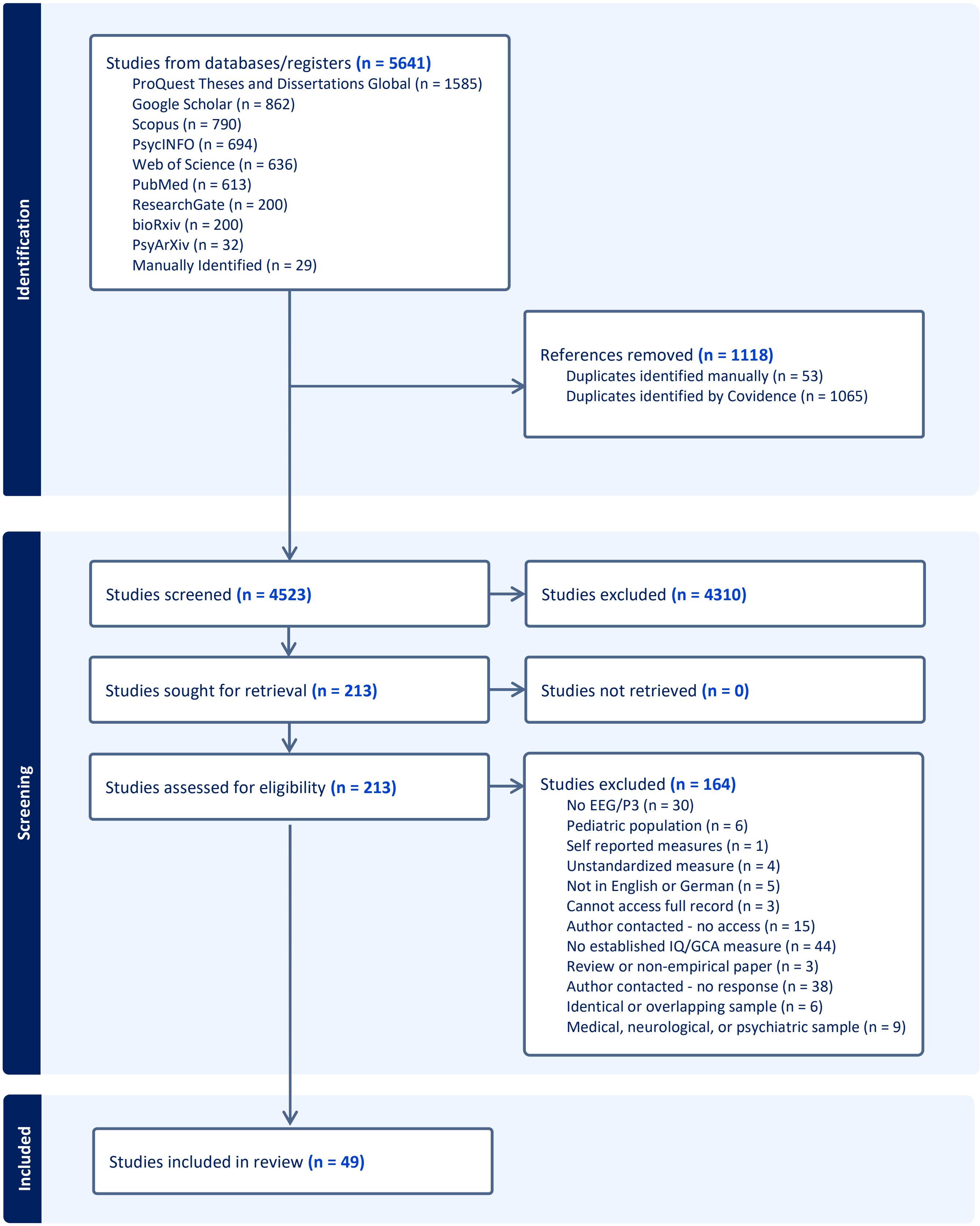

### Fig. 2

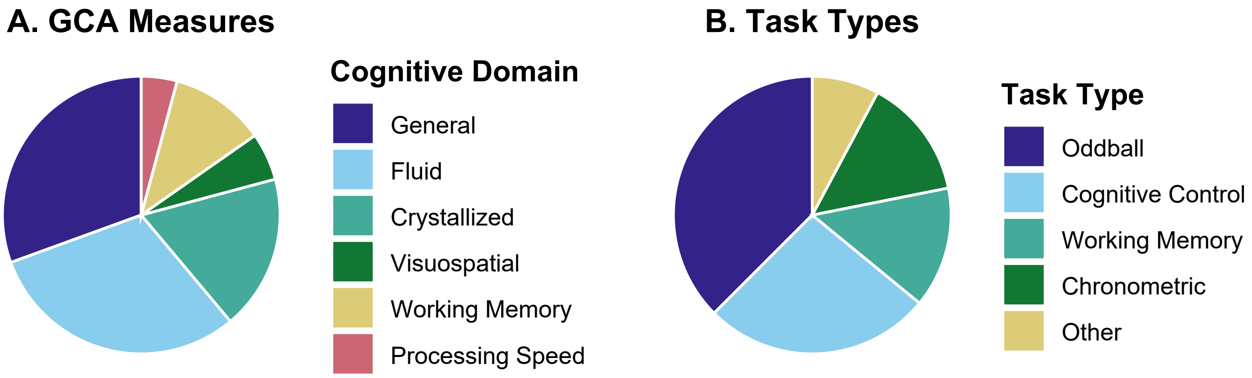

### Fig. 4

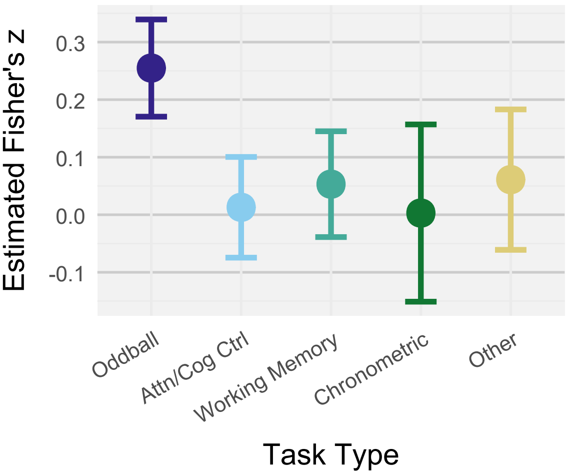

### Fig. 5

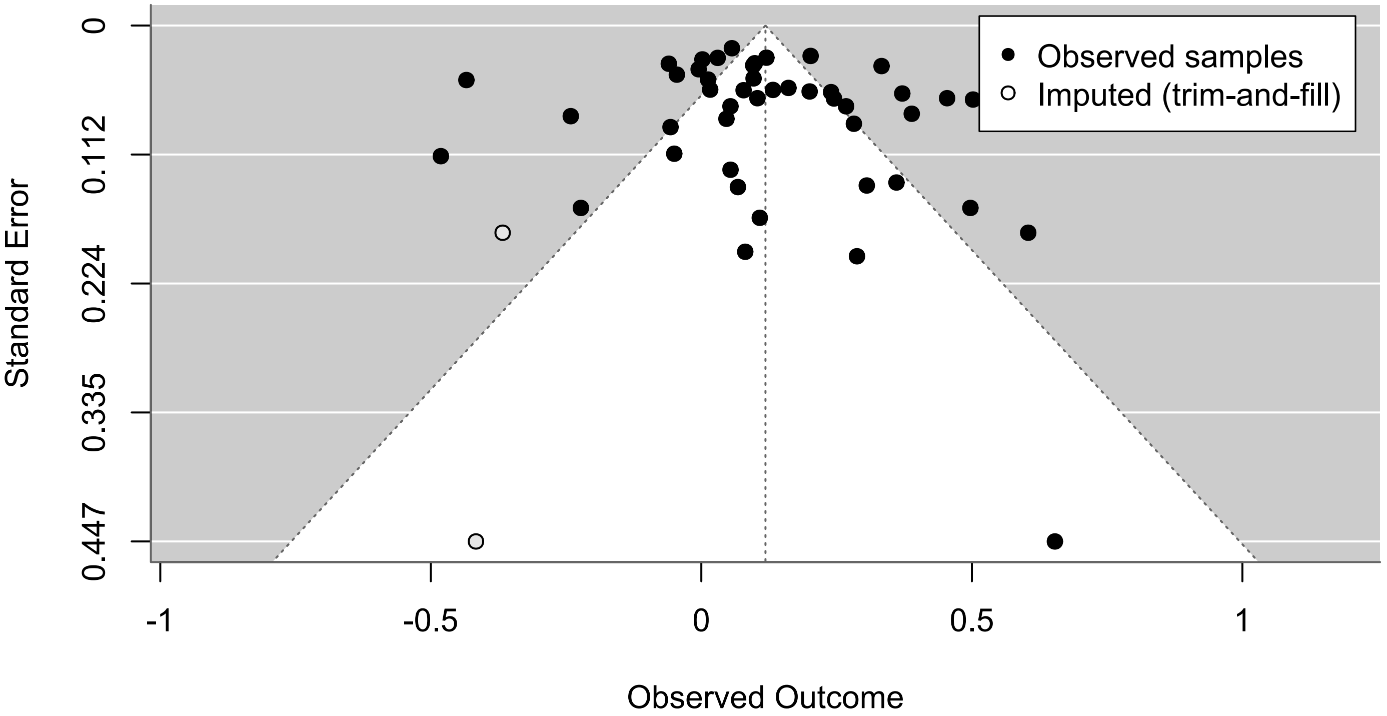

### Fig. 6

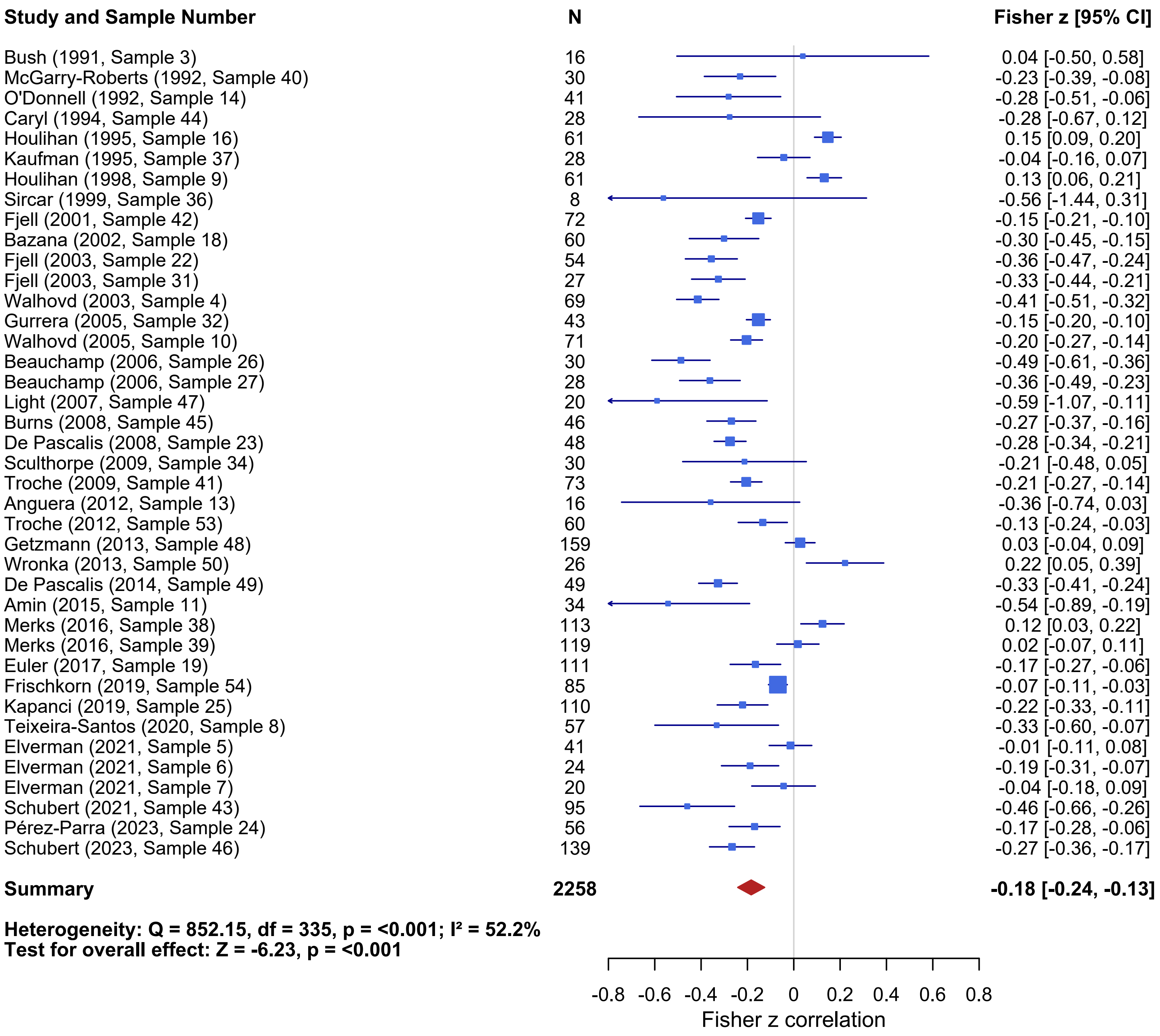

### Fig. 7

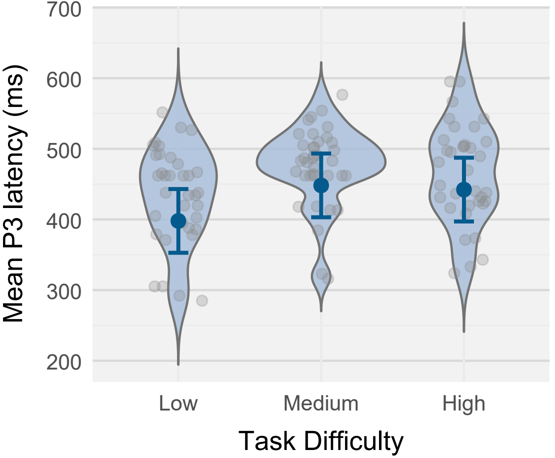

### Fig. 8

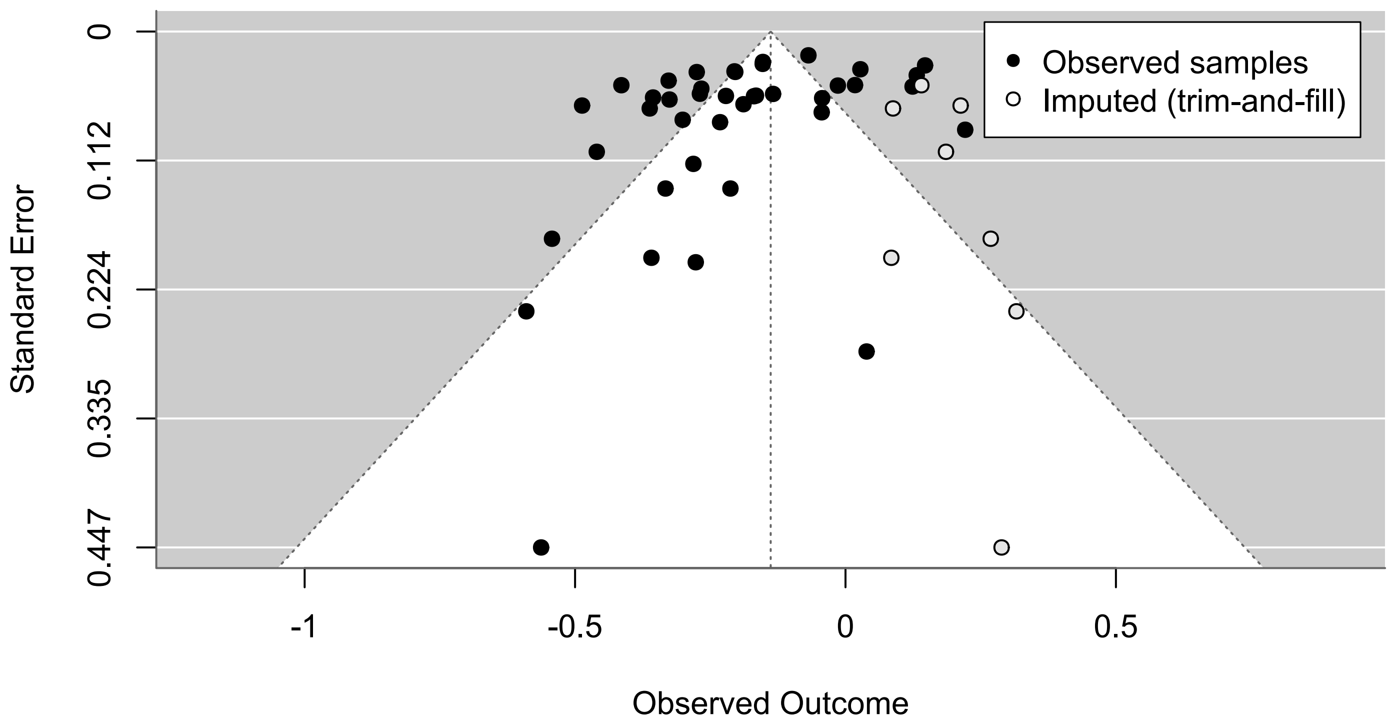
