## Supplementary material for "Human Cognitive Ability and the P300 Event-Related Brain Potential: A Systematic Review and Meta-Analysis": Fig. 3 (from another version of this preprint: version 1, 2026-02-16)

| Study / Sample | N | Fisher z [95% CI] |
|---|---|---|
| McGarry-Roberts (1992, Sample 40) | 30 | -0.24 [-0.40, -0.09] |
| Caryl (1994, Sample 44) | 28 | 0.29 [-0.10, 0.68] |
| Houlihan (1995, Sample 16) | 61 | 0.00 [-0.06, 0.06] |
| Kaufman (1995, Sample 37) | 28 | 0.24 [0.13, 0.35] |
| Alcorn (1996, Sample 20) | 49 | 0.12 [0.07, 0.17] |
| Katsanis (1997, Sample 33) | 128 | 0.25 [0.12, 0.37] |
| Houlihan (1998, Sample 9) | 61 | -0.00 [-0.08, 0.07] |
| Sircar (1999, Sample 36) | 8 | 0.65 [-0.22, 1.53] |
| Fjell (2001, Sample 42) | 72 | 0.03 [-0.02, 0.09] |
| Bazana (2002, Sample 18) | 60 | 0.39 [0.24, 0.54] |
| Fjell (2003, Sample 22) | 54 | 0.20 [0.09, 0.31] |
| Fjell (2003, Sample 31) | 27 | 0.37 [0.26, 0.49] |
| Walhovd (2003, Sample 4) | 71 | 0.10 [0.01, 0.19] |
| Gurrera (2005, Sample 32) | 43 | 0.20 [0.15, 0.25] |
| Beauchamp (2006, Sample 26) | 30 | 0.50 [0.38, 0.63] |
| Beauchamp (2006, Sample 27) | 28 | 0.53 [0.40, 0.66] |
| Light (2007, Sample 47) | 20 | 0.07 [-0.21, 0.34] |
| Burns (2008, Sample 45) | 46 | -0.06 [-0.23, 0.12] |
| De Pascalis (2008, Sample 23) | 48 | 0.33 [0.26, 0.40] |
| Russo (2008, Sample 17) | 45 | 0.45 [0.33, 0.58] |
| Sculthorpe (2009, Sample 34) | 30 | 0.36 [0.09, 0.63] |
| Troche (2009, Sample 41) | 73 | 0.10 [0.03, 0.16] |
| Anguera (2012, Sample 13) | 16 | 0.08 [-0.30, 0.47] |
| Troche (2012, Sample 53) | 60 | 0.16 [0.06, 0.27] |
| Getzmann (2013, Sample 48) | 159 | 0.10 [0.03, 0.16] |
| Wronka (2013, Sample 50) | 26 | 0.28 [0.12, 0.45] |
| De Pascalis (2014, Sample 49) | 49 | -0.05 [-0.13, 0.04] |
| Ramchurn (2014, Sample 55) | 16 | -0.48 [-0.70, -0.26] |
| Amin (2015, Sample 11) | 34 | 0.60 [0.25, 0.96] |
| Euler (2015, Sample 52) | 29 | 0.31 [0.03, 0.58] |
| Euler (2017, Sample 19) | 111 | 0.02 [-0.09, 0.13] |
| Varriale (2018, Sample 21) | 37 | 0.27 [0.13, 0.40] |
| Wongupparaj (2018, Sample 35) | 115 | -0.43 [-0.53, -0.34] |
| Zhang (2018, Sample 28) | 23 | -0.22 [-0.53, 0.09] |
| Zhang (2018, Sample 29) | 21 | 0.11 [-0.22, 0.43] |
| Zhang (2018, Sample 30) | 23 | 0.50 [0.19, 0.81] |
| Kapanci (2019, Sample 25) | 110 | 0.13 [0.02, 0.24] |
| Spironelli (2020, Sample 1) | 35 | 0.05 [-0.19, 0.30] |
| Elverman (2021, Sample 5) | 41 | 0.01 [-0.08, 0.10] |
| Elverman (2021, Sample 6) | 24 | 0.10 [-0.02, 0.23] |
| Elverman (2021, Sample 7) | 20 | 0.05 [-0.08, 0.19] |
| Li (2021, Sample 12) | 41 | -0.06 [-0.13, 0.00] |
| Löffler (2021, Sample 15) | 84 | -0.05 [-0.27, 0.17] |
| Markiewicz (2022, Sample 2) | 54 | 0.05 [-0.11, 0.20] |
| Kührt (2023, Sample 51) | 147 | 0.06 [0.02, 0.10] |
| Pérez-Parra (2023, Sample 24) | 56 | 0.08 [-0.03, 0.19] |
| **Summary** | **2371** | **0.13 [0.06, 0.19]** |

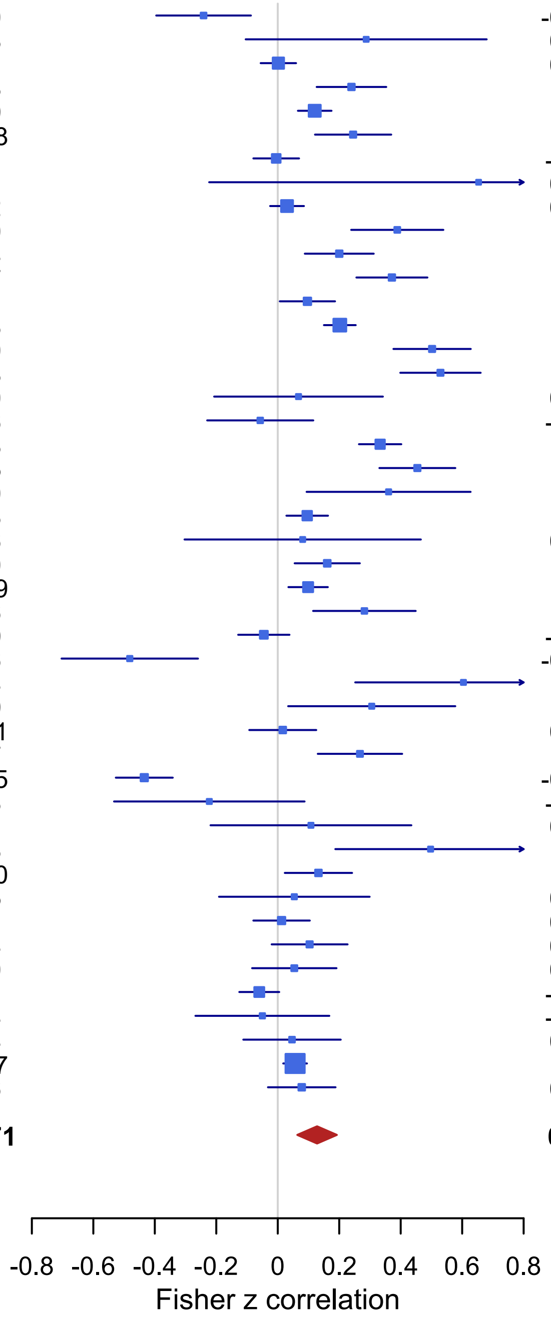


**Heterogeneity: Q = 914.8, df = 380, p = <0.001; I² = 60.36%**
**Test for overall effect: Z = 3.9, p = <0.001**
